# The SNARE Sec22b regulates phagosome maturation by promoting ORP8-mediated PI(4)P exchange at ER-phagosome contact sites

**DOI:** 10.1101/2022.04.12.487993

**Authors:** Nina Criado Santos, Samuel Bouvet, Flavien Bermont, Cyril Castelbou, Farah Mansour, Maral Azam, Francesca Giordano, Paula Nunes-Hasler

## Abstract

The precise control of phagosome maturation is critical for innate and adaptive immunity, determining whether phagocytosed material is destroyed or used to present antigens. We observed previously that non-fusogenic contacts between the endoplasmic reticulum (ER) and phagosomes, called membrane contact sites (MCS), are tethered by the calcium regulator STIM1 and fine-tune phagosomal maturation. The secretory pathway SNARE protein Sec22b has been implicated in controlling phagocytosis, phagosome maturation and antigen presentation, though its effects are controversial, and its mechanism of action poorly understood. Recently, Sec22b was shown to tether MCS at the plasma membrane without mediating membrane fusion. Here, we show that Sec22b localizes to and regulates the frequency of ER-phagosome contacts independently of STIM proteins. Sec22b knockdown and overexpression of a an MCS-disrupting mutant Sec22b-P33 induced only mild or no effect on global and local calcium signalling. However, Sec22b knockdown altered phagosomal phospholipids including PI(3)P, PI(4)P and PS, but not PI(4,5)P2. Increased PI(4)P in shSec22b cells was rescued by re-expression of Sec22b or the artificial MCS tether MAPPER but not the P33 mutant. Moreover, Sec22b co-precipitated and was co-recruited to phagosomes with the PS/PI(4)P lipid exchange protein ORP8. Expression of wild-type, but not mutant ORP8, also rescued phagosomal PI(4)P. Concordantly, Sec22b, MAPPER and ORP8 but not P33 or the ORP8 mutant decreased phagolysosome fusion in shSec22b cells. These results clarify a novel mechanism through which Sec22b controls phagosome maturation and beg a reassessment of the relative contribution of Sec22b-mediated fusion versus tethering to phagosome biology.

## Introduction

Phagocytosis is a critical immune process through which phagocytic immune cells such as neutrophils, macrophages and dendritic cells (DCs) engulf foreign particles into a membrane-enclosed vacuole. The ingested material is either destroyed or processed to antigens, rendering phagocytosis important to both innate and adaptive immunity. Phagosomes are formed by actin-driven membrane remodelling, followed by pinching off and sequential maturation involving fusion with endosomes and lysosomes which impart the phagosome with increasing degradative capabilities ^1, 2^. Endoplasmic reticulum (ER) membranes are also recruited to phagosomes, although the underlying mechanisms and their functional role has been debated ^3, 4^. In our previous research we have found that the ER-resident calcium (Ca^2+^) regulator STIM1 drives ER recruitment to phagosomes in neutrophils, dendritic cells and phagocytic fibroblast models to form structures comprised of tightly tethered (10-30 nm) associations with the phagosomal membrane called membrane contact sites (MCS) ^5–9^. We found that MCS fostered localized calcium signals that promoted actin disassembly and lysosome fusion, driving phagocytic ingestion rates and maturation. However, even in *Stim1^-/-^Stim2^-/-^* double knock-out cells, considerable STIM-protein independent MCS remained ^6^, suggesting additional MCS tethers may regulate phagocytosis.

Sec22b is a multifunctional protein of the Soluble N-ethylmaleimide-Sensitive Factor Attachment Proteins Receptor (SNARE) family generally associated with the function of mediating membrane fusion, that localizes to both the ER and ER-to-Golgi intermediate compartment (ERGIC) ^10, 11^. Sec22b was originally described in the regulation of both retrograde and anterograde trafficking in the secretory pathway, in a role partially redundant with that of closely related SNARE YKT6 ^10, 12, 13^. In addition to a short cytoplasmic C-terminus, single transmembrane domain, and SNARE motif, it also possesses a large N-terminal longin domain that regulates its localization and function ^11, 14, 15^. Notably, it was shown to tether the ER to the plasma membrane (PM) in bona-fide MCS without mediating fusion, due to its longin domain-mediated exclusion of SNAP-25 from SNARE complexes, in neurons and Hela cells ^15, 16^. In macrophages, both positive and negative roles for Sec22b in controlling phagocytic rates were observed ^17, 18^, whereas in DCs Sec22b knockdown promoted phagosome maturation, increasing phago-lysosome fusion without affecting phagocytic rates ^19^. In all cases Sec22b was proposed to achieve these effects by mediating fusion of either ER or ERGIC membranes with the phagosome, though fusion per se was never assessed directly. In addition, it is unclear how fusion of ER or ERGIC could reduce phagosome fusion with lysosomes.

In light of our observations of the intimate but non-fusogenic association of ER membranes with phagosomes in the context of STIM1-mediated MCS, and studies showing the non-fusogenic tethering role of Sec22b at the PM, we sought to determine whether Sec22b controls phagocytosis or phagosome maturation through a tethering role at ER-phagosome contacts. To this end we utilized a fibroblast cell line rendered phagocytic by ectopic expression of the IgG receptor FCGR2A, a cell model that is much more amenable to transfection of multiple fluorescent constructs and high-resolution imaging than natural phagocytes, and which we have extensively characterized and shown to recapitulate many key aspects of the phagocytic process ^5, 6^. By overexpressing and knocking down Sec22b in the presence and absence of STIM1, we show that Sec22b is present at and regulates ER-phagosome MCS independently of STIM proteins. Since modulation of Sec22b expression imparted only mild, both positive and negative roles on Ca^2+^ signalling, we further investigated whether Sec22b might instead regulate non-vesicular lipid transfer at these contacts, since it is another function often associated with MCS that Sec22b was shown to regulate at the PM ^8, 9, 15, 16^. We provide evidence that Sec22b regulates phagosomal levels of phospholipids at least in part by recruiting the lipid transfer protein ORP8, and that this in turn controls phago-lysosome fusion.

## Results

### Sec22b localizes to and modulates ER-Phagosome MCS

Given the fact that Sec22b operates as a tether between the ER and PM, we first investigated whether Sec22b localizes to ER-phagosome contact sites (ER-Phg MCS). Sec22b has been reported to localize to both the ER and ERGIC in various cell types including neurons and DCs, with an estimated 50:50 distribution in yeast and HEK293T cells, and a visibly greater proportion in the ERGIC in other cells ^16, 19–21^. Mouse embryonic fibroblasts (MEFs) rendered phagocytic by overexpression of myc-FCGR2A were used here as a phagocytic model that is genetically tractable and whose ER-Phg MCS we have characterized extensively. Immunostainings with two commercial antibodies against Sec22b (SC: Santa Cruz Biotechnologies sc-101267 and SYSY: Synaptic Systems 186003) were performed in phagocytic MEFs exposed to IgG-coated targets. Both antibodies showed staining of variable quality, but consistent with both ER and ERGIC localization, although the SC antibody showed a larger ER pool, while the SYSY antibody preferentially highlighted ERGIC structures (Fig. 1A-C). Nevertheless, even when employing the SYSY antibody, immunostaining of cells co-transfected with the ERGIC marker GFP-ERGIC-53 (Fig. 1A) or both GFP-ERGIC-53 and KDEL-tagged, ER-targeted-RFP (RFP-KDEL) (Fig. 1D) displayed punctate periphagosomal Sec22b accumulations reminiscent of contact sites that did not co-localize with the ERGIC marker (Fig. 1A, Site 1), in addition to areas that did (Fig. 1A, Site 2). A similar pattern was observed when co-expressing mCherry-Sec22b and GFP-ERGIC-53 although a larger ER pool was evident using the overexpressed protein, similar to the SC antibody (Fig. 1B). Co-expression of mCherry-Sec22b with FCGR2A-GFP, or EGFP-Sec22b with myc-FCGR2A highlighted the difference between the continuous staining of a protein present on the phagosomal membrane (the Fc-receptor) and the punctate nature of the Sec22b periphagosomal localization (Fig. 1E, Fig. S1A).

**Figure 1.**
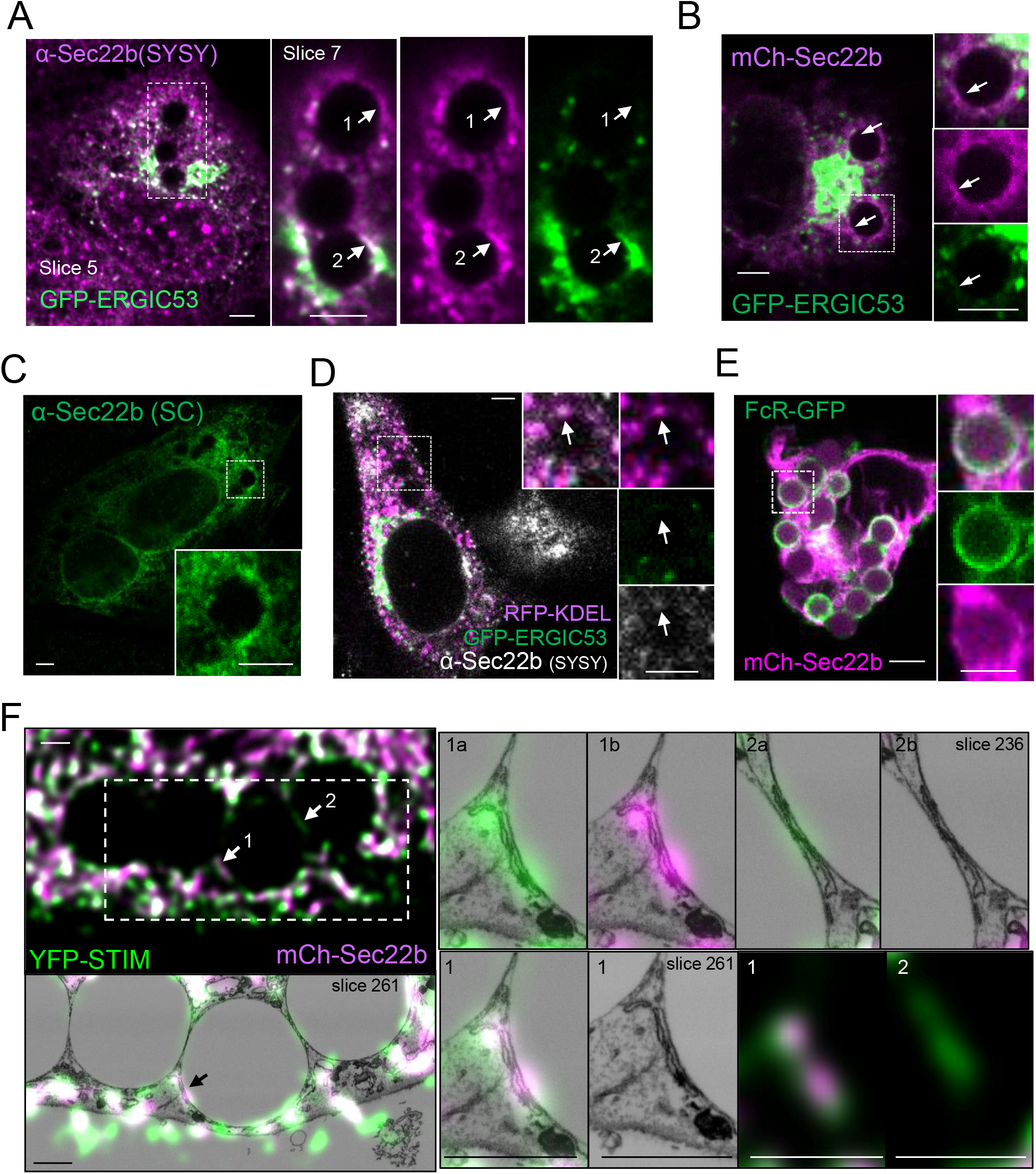
Sec22b localizes to ER-phagosome membrane contact sites. **A-B**. Confocal images of Immunostaining of endogenous Sec22b using the Synaptic Systems (SYSY) antibody (**A**, magenta) or fluorescence of mCherry (mCh)-Sec22b (**B**, magenta) in phagocytic MEFs overexpressing GFP-ERGIC-53 (green) and exposed to IgG-beads for 30 min. Sec22b periphagosomal accumulation containing (arrow 1) and devoid (arrow 2 in A, and arrow in B) of the ERGIC marker. Slice 5 of the total stack is shown to highlight total cell staining while Slice 7 is shown in the inset to highlight periphagosomal accumulation. **C**) Immunostaining of endogenous Sec22b (green) in phagocytic MEFs exposed to IgG-beads for 30 min with the Santa Cruz antibody (SC). **D**) Immunostaining of endogenous Sec22b (white) in phagocytic MEFs overexpressing both the ER-marker RFP-KDEL (magenta) and GFP-ERGIC-53 (green), exposed to IgG-beads for 30 min. Arrow: periphagosomal accumulations of ER marker that are devoid of ERGIC marker and contain a faint but visible Sec22b signal. **E**) MEFs overexpressing FCGR2A-GFP (FcR, green) and mCh-Sec22b (magenta) and exposed to IgG-beads for 15 min. **F)** 3D-CLEM analysis of phagocytic *Stim1^-/-^; Stim2^-/-^* (DKO) MEFs overexpressing mCh-Sec22b (magenta) and YFP-STIM1 (green) exposed to IgG-beads for 30 min. Arrow 1: periphagosomal accumulation of Sec22b co-localizing with STIM1 in an MCS structure. Arrow 2: MCS containing only STIM1 but not Sec22b. One deconvolved confocal slice corresponds approximately to 75 EM slices. Slice 261 of the EM stack is shown in the large images and insets of site 1, while slice 236 is shown for site 2. Insets labelled “a” show an overlay of the EM with the green channel only, those labelled “b” with the red channel only. For A-E bars = 3 µm, for F bars = 1 µm.

We have previously shown that the ER-resident Ca^2+^ regulator STIM1 localizes to ER-Phg MCS ^5, 7^. To visualize bona-fide ER-Phg MCS, phagocytic *Stim1^-/-^/Stim2^-/-^* double knock-out MEFs (DKO) that were co-transfected with YFP-STIM1 and mCherry-Sec22b were exposed to IgG-coated targets and analysed by 3D correlation light electron microscopy (3D-CLEM). DKO cells were employed to mitigate the effects of STIM1 overexpression ^22, 23^. Periphagosomal Sec22b puncta co-localized extensively with STIM1 (Fig. 1F, Site 1). Analyses of these areas of co-localization using 3D-CLEM not only confirmed that these structures did indeed correspond to bona-fide MCS, it also allowed a complete inspection of structures above and below the MCS that could contribute to a co-localization signal. In the structures indicated by the arrows (Fig. 1F, Site 1; Fig. S1B) only other MCS and no other vesicular structures that could correspond to endosomes or ERGIC vesicles were present above or below the ER contacting the phagosome (Fig. S1B), strongly suggesting that Sec22b within the ER was indeed localizing to ER-Phg MCS. While most (though not all) periphagosomal structures containing Sec22b also contained STIM1, some periphagosomal MCS marked by STIM1 that were devoid of Sec22b were captured in the CLEM image (Fig. 1F, Site 2). This indicates that inclusion in an ER-Phg MCS is selective and not an automatic consequence of Sec22b’s overexpression within the ER, or its co-expression with STIM1, whose overexpression causes an increase in the frequency and size of MCS ^5, 23^. Furthermore, it suggests that similar to ER-PM MCS, phagosomal MCS are also heterogeneous and may have a variable composition that differs not only from bulk ER, but also between different contacts ^8, 9, 24, 25^.

To determine whether Sec22b contributes to ER-Phg MCS tethering, an assessment of the effect of Sec22b knockdown on the frequency of periphagosomal MCS was performed. Stable cell lines were established by lentiviral transduction of control (shCTR) or short hairpin RNA (shRNA) targeting Sec22b (shSec22b) followed by antibiotic selection. A knockdown efficiency of ⁓80% was confirmed by Western blot (Fig. 2A, Fig. S2A-B). Cells were then exposed to IgG-coupled targets, fixed and embedded for transmission electron microscopy (TEM) ^5^. Quantification of periphagosomal MCS by morphological assessment of random TEM slices was performed as previously described ^5^. A ⁓30% loss in the frequency of periphagosomal MCS was observed, from an average of 4.5 contacts per phagosome in shCTR cells to 3.0 in shSec22b cells, indicating that Sec22b contributes to ER-Phg MCS tethering (Fig. 2B-C). Whereas the median size of individual contacts was slightly lower (93 vs 81 nm in shCTR vs shSec22b, Fig. 2C), a small population (3%) of very large (>400nm) contacts were observed uniquely in shSec22b cells (Fig. 2C, red bracket) resulting in an average size that was not significantly different. Recruitment of Sec22b to ER-Phg contact sites was independent of STIM protein expression since Sec22b still localized to ER-Phg MCS, as indicated by CLEM analysis, in phagocytic *Stim1^-/-^/Stim2^-/-^* (DKO) cells co-expressing mCherry-Sec22b and the artificial MCS marker EGFP-MAPPER which labels MCS but does not majorly affect Ca^2+^ signalling ^26, 27^. Interestingly, in contrast to STIM1, Sec22b and MAPPER co-localization was less pronounced and appeared to segregate to adjacent MCS zones (Fig. 2D, sites 1 and 2). MCS devoid of either Sec22b or MAPPER were also observed (Fig. 2D, Site 3), again highlighting the heterogeneity of ER-Phg MCS. In addition, loss of Sec22b in DKO cells led to a loss of MAPPER recruitment to phagosomes (Fig. 2E), further indicating that Sec22b contributes to ER-Phg MCS tethering independently of STIM proteins. Collectively, these data demonstrate that ER-bound Sec22b localizes to and influences the frequency of ER-Phg MCS, supporting the view that ER-localized Sec22b may play an additional role during phagocytosis.

**Figure 2.**
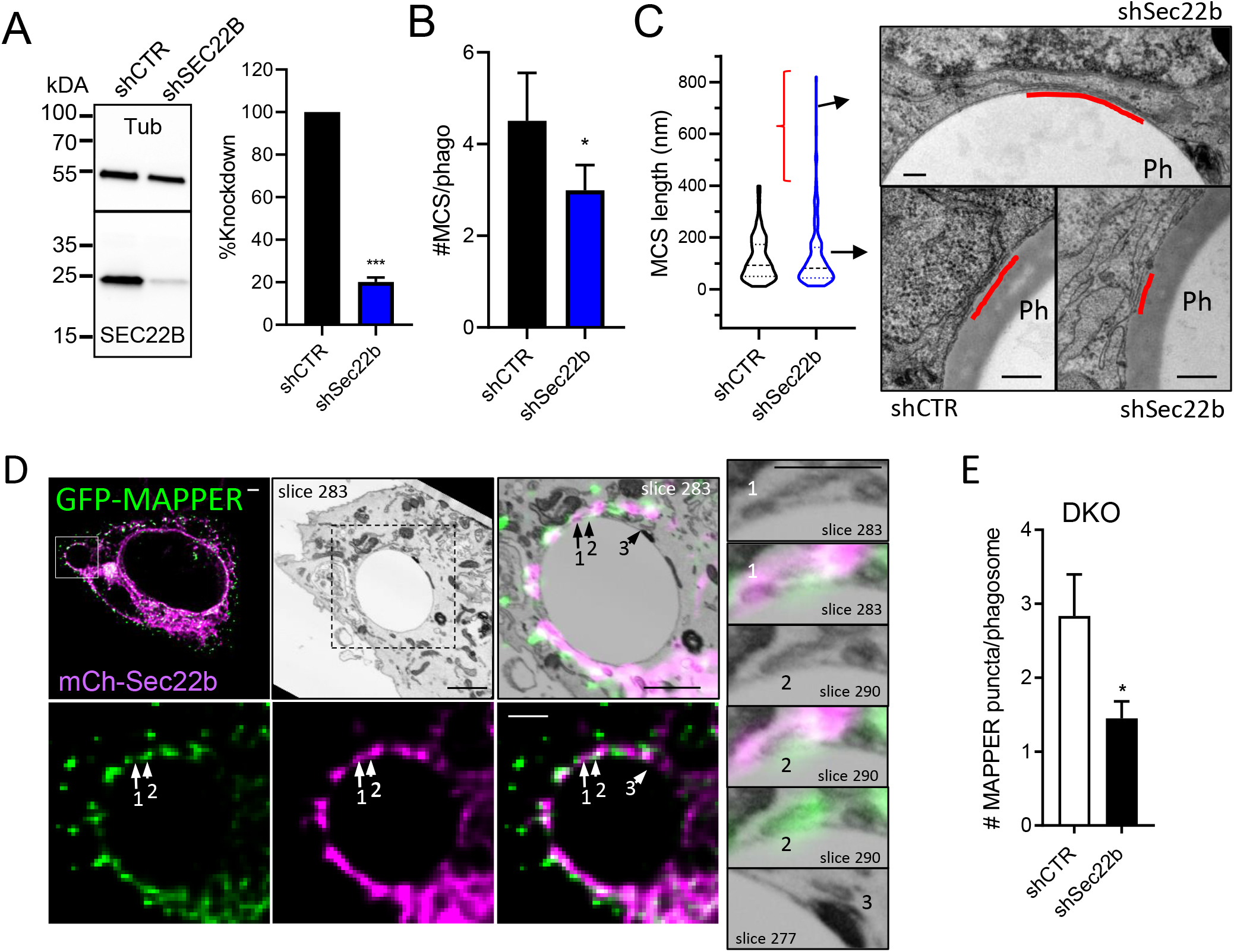
Sec22b regulates phagosomal contact sites independently of STIM1. **A)** Western blot of stable MEF cell lines expressing control (shCTR) or shRNA directed against Sec22b (shSec22b) (n=12, blots normalized to anti-tubulin controls, expressed relative to shCTR, see also Fig. S2A-B). **B)** Quantification of periphagosomal MCS frequency (number of MCS <30 nm to phagosomal membrane per phagosome) in transmission EM slices (n=3, shCTR/shSec22b). **C)** Quantification of MCS length (left panel) revealed a greater diversity in MCS length in shSec22b cells (F test comparing variances p<0.0001), with a small (3%) population of very large contacts that were not present in shCTR cells (red bracket, example in top right panel image). Median MCS length was smaller in shSec22b (93.05/81.90 nm, typical examples in bottom right panel images) but mean values were similar. (n=3, 172/145 contacts in 44/45 phagosomes shCTR/shSec22b). Bars = 100 nm. **D)**. 3D-CLEM analysis of STIM-null DKO phagocytic MEFs overexpressing GFP-MAPPER (green) and mCh-Sec22b (magenta) and exposed to IgG-beads for 30 min. Arrow 1: Sec22b-positive MCS (slice 283, upper insets). Arrow 2: MAPPER-positive MCS (slice 290, lower insets), Arrow 3: MCS devoid of both markers (slice 277, bottom inset). Bars = 1 µm. **E)** Quantification of periphagosomal GFP-MAPPER puncta in DKO cells stably transfected with shCTR or shSec22b show a loss of MAPPER recruitment to phagosomes upon Sec22b knockdown (n=3/5 coverslips in shCTR/shSec22b). Bars graphs show means +SEM. * p< 0.05, *** p<0.001.

### Sec22b is not required but can modulate Ca^2+^ signalling

The Ca^2+^ regulators STIM1 and its close homolog STIM2 function both at the ER-PM MCS, promoting global Ca^2+^ signals, as well as at ER-Phg MCS, where particularly STIM1 plays a major role in generating localized Ca^2+^ signals or hotspots ^5, 6^. Given the extensive colocalization of STIM1 and Sec22b, we next investigated whether Sec22b cooperates with STIM1 at MCS to regulate Ca^2+^ signalling, starting with ER-PM MCS which are easier to image. Total internal reflection microscopy (TIRF) analysis allowed simultaneous imaging of Sec22b and STIM1 recruitment to ER-PM MCS, visible as puncta at the PM plane, during the activation of store-operated Ca^2+^ entry (SOCE) (Fig. 3A). Targeted activation of SOCE that does not rely on receptor-mediated or lipid signalling is achieved by applying the sarcoplasmic reticulum ATPase inhibitor thapsigargin (Tg) which releases Ca^2+^ stored within the ER. This event promotes STIM1 activation, oligomerization and translocation to ER-PM MCS, steps which are necessary for STIM1 interaction with partner channels of the ORAI and TRPC families, initiating Ca^2+^ influx ^28^. In order to determine whether Sec22b could influence STIM1 recruitment to ER-PM MCS, Tg was applied to either WT or DKO MEFs overexpressing YFP-STIM1 and mCherry-Sec22b or RFP-KDEL as control, in the absence of external Ca^2+^ (which prevents the negative feedback of STIM1 de-activation upon Ca^2+^ influx) ^29^. As above, DKO cells were employed to mitigate the effects of STIM1 overexpression, which when excessive can be saturating and inhibitory to Ca^2+^ signalling ^22, 28, 30^. TIRF plane puncta, representing ER-PM MCS, increased in numbers and size following a sigmoidal time function (Fig. 3A-B, Fig. S2C). Puncta were larger in both WT and DKO cells (Fig. 3A-C) and more numerous (Fig. S2C) in WT cells overexpressing Sec22b as compared to KDEL controls. In addition, STIM1 recruitment was accelerated, manifesting as either a decreased slope parameter in the sigmoidal curve fit in WT cells, or a decreased time constant (t_50%_) in DKO cells indicating that Sec22b expression facilitates STIM1 translocation to ER-PM MCS (Fig. 3C, Fig. S2C). On the other hand, Sec22b was not required for STIM1 recruitment to ER-PM MCS as Sec22b knockdown had no significant effect on the kinetics of YFP-STIM1 recruitment (Fig. 3D, Fig. S2D). Next, whether the Sec22-dependent changes in STIM1 recruitment have an impact on global Ca^2+^ influx was assessed. To this end a classical assay for assessing SOCE ^22, 28, 30^ was employed where cells loaded with the ratiometric Ca^2+^-sensitive dye Fura-2 are exposed to Tg in Ca^2+^-free medium as above, and the maximum slope and peak amplitude of the Fura-2 Ca^2+^ signal are measured upon the re-addition of extracellular Ca^2+^. Surprisingly, despite the effect on STIM1 recruitment observed by TIRF, SOCE was unchanged in WT cells either overexpressing mCh-Sec22b in combination with YFP-STIM1, or expressing EGFP-Sec22b alone, as compared to RFP or EGFP-KDEL controls respectively (Fig. 3E-F). This was in contrast to overexpression of the Sec22b-P33 mutant, a construct where insertion of a 33 amino-acid proline tract was show to extend the MCS gap distance, compromising Sec22b’s MCS function without affecting secretory transport ^16^. In this case, global signals were faster and larger (Fig. 3F), indicating that MCS tethering by wild-type Sec22b may have both activating and inhibitory effects on the SOCE machinery that cancel out, but are revealed upon Sec22b mutation. In support of this hypothesis is the observation that in contrast to overexpression, a small but significant increase in the peak amplitude was observed upon Sec22b knockdown (Fig. 3G). This effect was not due to a compensatory increase in STIM1 expression upon Sec22b knockdown, as STIM1 levels were unchanged in shSec22b MEF WT cells as compared to shCTR (Fig. 3H, Fig. S2B). Nevertheless, this effect was still dependent on STIM1 since it was abolished in *Stim1^-/-^* single knock-out (SKO) cells, which retain measurable residual influx (Fig. 3F), when these cells were transfected with shCTR or shSec22b (Fig. 3I, Fig. S2B).

**Figure 3.**
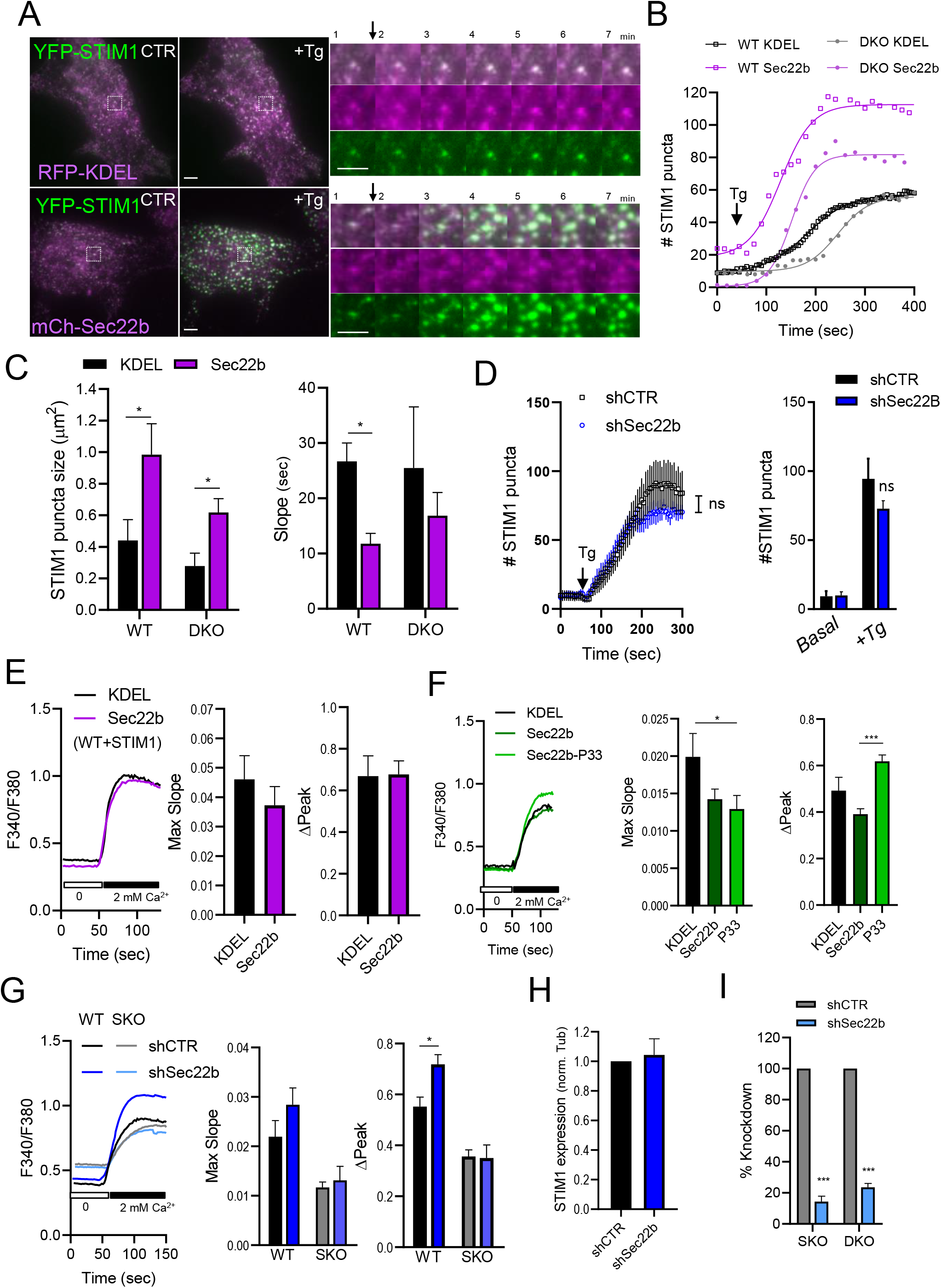
Sec22b plays both positive and negative roles in calcium signalling. **A-C)** TIRF analysis of wild-type (WT) and DKO MEFs transfected with YFP-STIM1 in combination with ER marker RFP-KDEL or mCh-Sec22b. Cells washed in calcium-free imaging buffer (CF) were imaged every 15 sec, and 1 µM thapsigargin (Tg) was added after 1 min. The number (**B,** See also Fig. S2C), size (**C,** left panel), and kinetics (**C,** right panel) of STIM1 puncta appearance at the TIRF plane, representing ER-plasma membrane (PM) MCS, were quantified 6 min after Tg addition. Larger STIM1 puncta appeared more rapidly upon Sec22b overexpression. In (**A**) error bars are omitted for clarity, mean values and Boltzman sigmoidal curve used to calculate slope parameters are shown. n=8/11;3/5 coverslips WT;DKO KDEL/Sec22b). **D)** Similar experiments as in A-C revealed Sec22b knockdown in DKO cells does not impact mCh-STIM1 recruitment to ER-PM MCS (n=13/12 shCTR/shSec22b, see also Fig. S2D). **E-F)** Global store-operated calcium influx was measured using classic Fura-2-based Tg-Calcium re-addition assay (see methods) in WT MEFs transfected with YFP-STIM1 and RFP-KDEL or mCh-Sec22b (**E**) or in WT MEFs transfected only with GFP-KDEL, EGFP-Sec22b, or EGFP-Sec22b-P33 MCS-disrupting mutant (**F**). Influx was faster compared to KDEL and higher compared to wild-type Sec22b in cells expressing the P33 mutant. Error bars are omitted from traces for clarity. **G**) Similar measurements of global calcium influx were performed in WT and *Stim1^-/-^* (SKO) cells. Peak influx was higher in WT shSec22b cells (n=15/13;19/20 WT;DKO shCTR/Sec22b). **H**) Quantification of Western blots of WT MEF shCTR and shSec22b lysates show no difference in STIM1 expression upon Sec22b knockdown (n=8, blots normalized to anti-tubulin controls, expressed as relative to shCTR, See also Fig. S2B). **I**) Quantification of Western blots of SKO and DKO MEF shCTR and shSec22b lysates show robust knockdown (n=5/5 SKO/DKO, blots normalized to anti-tubulin controls, expressed relative to shCTR, see also Fig. S2A-B). Bar graphs show means +SEM, *p<0.05, *** p<0.001.

We then turned our attention to assessing whether Sec22b could impact localized Ca^2+^ signalling during phagocytosis, which we hypothesized might be reduced upon loss of Sec22b expression in light of the decreased frequency of phagosomal MCS (Fig. 2B). To this end we employed a previously established Ca^2+^ hotspot assay based on the Ca^2+^-sensitive dye Fluo-8 where we have observed that about 50% of hotspots were dependent upon STIM1-mediated opening of phagosomal Ca^2+^ channels, with ER Ca^2+^ release through IP_3_R contributing to the majority of the remaining signal ^5, 6^. Phagocytic shCTR or shSec22b Fluo-8-loaded WT MEFs were thus exposed to IgG-coated targets and the frequency of periphagosomal Ca^2+^ hotspots within a 750 nm distance from the phagosomal border were quantified after 30 min of ingestion. However, hotspot frequency was unchanged (Fig. 4A) and phagocytic index was similar under these conditions (Fig. 4B). This indicates that Sec22b-dependent MCS may not be geared towards Ca^2+^ signalling, though perhaps a cancelling out of the reduced MCS but increased Ca^2+^ influx at remaining MCS could also explain these results. Taken together these results suggest a complex and probably indirect relationship between Sec22b and Ca^2+^ signalling. Sec22b is not required for global or localized Ca^2+^ signalling, but rather, depending on the context it can have both positive and negative modulatory roles that maybe secondary, i.e. through its influence on the shape and composition of MCS. These observations nevertheless further strengthen the idea that Sec22b localizes to phagosomal MCS and warrant further investigation on how this tethering role could impact phagocytic function.

**Figure 4.**
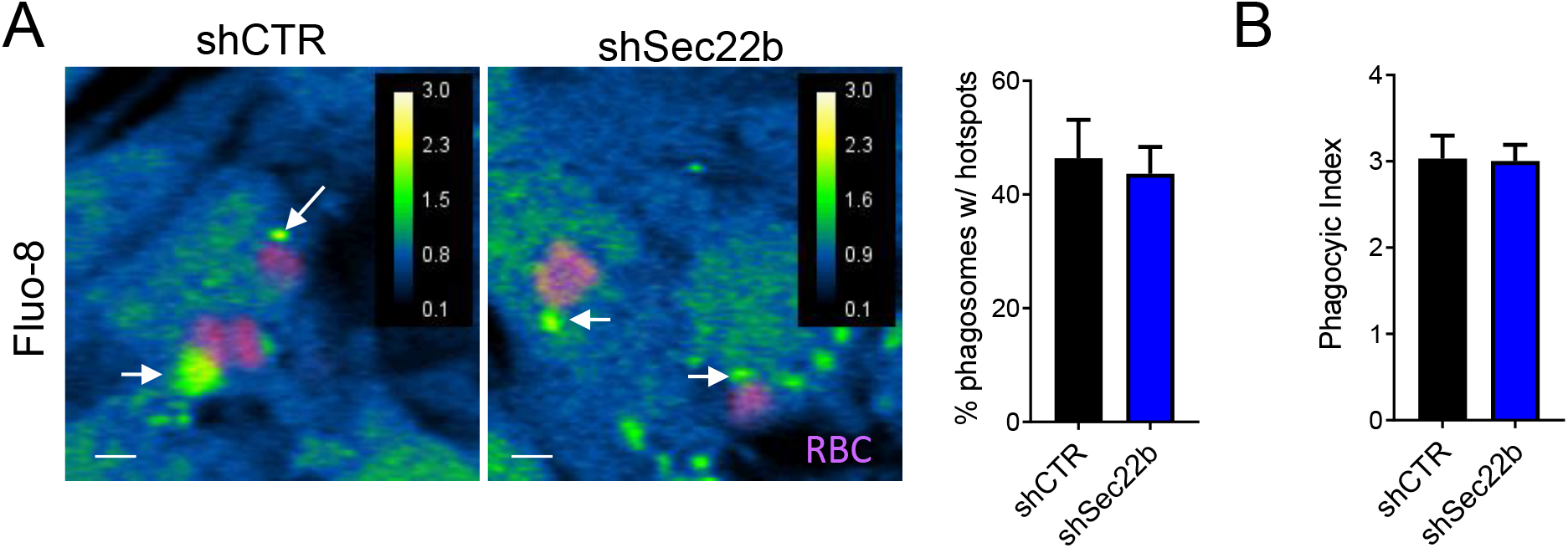
Sec22b knockdown does not affect periphagosomal calcium hotspots. **A-B)** Periphagosomal calcium hotspots(A) and phagocytic index (B) were quantified in shCTR and shSec22b phagocytic WT MEFs loaded with Fluo-8 (blue-green ratio pseudocolor) cells exposed to IgG-RBC (magenta) for 30 min. Sec22b knockdown did not impact the frequency of hotspots or phagocytic index. (n=7/8 shCTR/shSec22b). White bar = 3 µm. Bar graphs show means +SEM.

### Sec22b-mediated MCS tethering regulates phagosomal phospholipids

One of the functions attributed to MCS is to allow non-vesicular lipid transfer ^8, 9^. The yeast homolog Sec22 interacts with non-vesicular lipid transfer proteins (LTPs) Osh2 and Osh3, and temperature sensitive mutations of Sec22 led to an abnormal accumulation of PM levels of phosphatidylinositide-4-phosphate (PI(4)P) ^16^. Thus, we postulated that since Sec22b-dependent MCS were not majorly involved in Ca^2+^ signalling they might be instead involved in regulating phagosomal PI(4)P levels by promoting non-vesicular lipid transfer at phagosomal MCS. To test this hypothesis we utilized the PI(4)P probe 2xP4M ^31^ which has previously been employed to investigate phagosomal PI(4)P levels ^32^. Phagocytic shCTR and shSec22b MEFs were co-transfected with 2xP4M-GFP and the phosphatidylinositide-3-phosphate (PI(3)P) probe tagRFP-FYVE ^33^, where the PI(3)P probe was intended to serve as control. Cells were exposed to IgG-coupled beads, and live confocal images recorded for 30-40 min. The phagosomal signal of the probes were normalized to the initial average total cell signal in order to control for differences in expression levels, and represent the phagosomal lipid enrichment over time. In shCTR cells a strong initial peak followed by a rapid decrease of phagosomal PI(4)P was observed, after which two phagosomal populations were apparent. In the majority, phagosomal PI(4)P remained low, and in a minority, phagosomes displayed some continuing fluctuations in PI(4)P, similar to macrophages ^32^ (Fig. 5A). Whereas the initial peak increase in PI(4)P was comparable to controls, the fraction of phagosomes with continuing PI(4)P fluctuation was increased such that the average level of phagosomal PI(4)P between 5-25 min after the initial peak was significantly higher in Sec22b-depleted cells (Fig. 5A-B). Surprisingly, PI(3)P phagosomal levels were also de-regulated by Sec22b depletion where a lower average phagosomal level was observed between 5 and 10 min after ingestion, but not at later time points (Fig. 5C). However, this phenotype did not extend to all phagosomal phosophoinositides as PI(4,5)P2 levels, measured using the PLCδ-PH domain probe ^34^ were similar between the shCTR and shSec22b cells (Fig. 5D). The effects on phagosomal PI(4)P were reproduced in phagocytic MEF cells expressing 2xP4M-GFP without co-expression of other probes, when Sec22b was transiently depleted using an siRNA pool containing sequences that were distinct from the shSec22b sequence and reduced Sec22b expression by 63% (Fig. 5E, Fig. S3A,D). These results argue against confounding off-target effects of the shRNA sequence or the co-expression of the PI(3)P probe. Importantly, re-expression of an shRNA-resistant (shR) EGFP-tagged wild-type Sec22b as compared to EGFP-KDEL in combination with 2xP4M-mCherry rescued the PI(4)P phenotype, whereas shR-Sec22b-P33 did not (Fig. 5F), despite robust periphagosomal recruitment (Fig. S3E). This suggests a direct involvement of Sec22b MCS tethering in phagosomal PI(4)P regulation. Moreover, increasing MCS tethering by overexpression of MAPPER also lead to the rescue of phagosomal PI(4)P levels in Sec22b-depleted cells (Fig. 5F), supporting the view that Sec22b regulates phagosomal lipids by promoting MCS tethering. At both the plasma membrane and Golgi PI(4)P levels are regulated by LTPs in a counter-exchange mechanism with phosphatidylserine (PS) or cholesterol respectively that is powered by the dephosphorylation of PI(4)P in the ER by Sac1 ^35–37^. Thus, whether PS is affected by Sec22b depletion was examined by overexpressing an RFP-tagged PS-binding probe Lact-C2 ^38^. Indeed, in Sec22b-depleted cells a small but still significant decrease in phagosomal PS levels was detected (Fig. 5G). Together these data are consistent with a putative bidirectional transfer of PI(4)P and PS across ER-Phg MCS.

**Figure 5.**
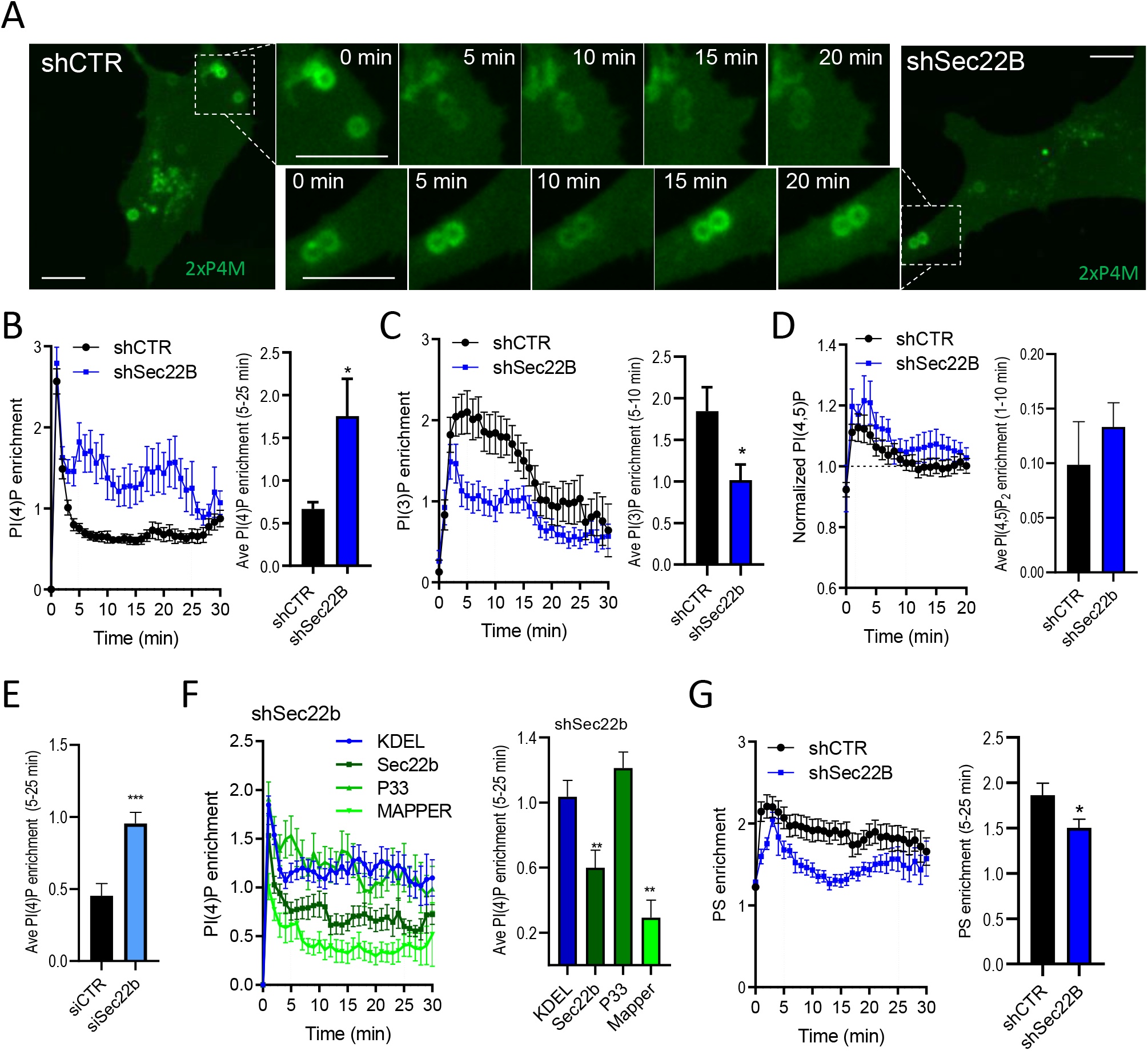
Sec22b controls the levels of phagosomal phospholipids. **A-C)** Live spinning-disk confocal microscopy of phagosomal PI(4)P and PI(3)P respectively, in shCTR and shSec22b phagocytic MEFs transfected with GFP-2xP4M **(A-B)** and TagRFP-FYVE(EE1A) **(C),** exposed to IgG-beads for 30-40 min, and imaged at a frequency of 1 frame/min. PI(4)P was significantly increased between 5-25 min and PI(3)P was decreased between 5-10 min after target internalization in shSec22b cells (n=8/10 coverslips shCTR/shSec22b). **D)** A similar analysis in cells expressing GFP-C1-PLCdelta-PH, showed no change in phagosomal levels of PI(4,5)P2 1-10 min after internalization. (n=8/6 coverslips shCTR/shSec22b) **E)** Increased phagosomal PI(4)P between 5-25 min after target internalization in WT phagocytic MEFs expressing GFP-2xP4M and transfected with siCTR or siSec22b (50 nM) (n=7/9 siCTR/siSec22b, see also Fig. S3A) **F)** Transfection of shSec22b phagocytic MEFs with mCh-2xP4M and either shR-EGFP-Sec22b or GFP-MAPPER reduced phagosomal PI(4)P levels as compared to GFP-KDEL controls, whereas shR-EGFP-Sec22b-P33 did not. (n= 19/14/5/6 KDEL/Sec22b/MAPPER/P33, see also Fig. S3E). **G)** Decreased phagosomal PS between 5-25 min after target internalization in shSec22b as compared to shCTR phagocytic MEFs transfected with mRFP-Lact-C2 (n=5/7 shCTR/shSec22b). White bars = 10 µm. Traces are means +/- SEM, bar graphs means + SEM. * p<0.05, **p<0.01, *** p<0.001.

### ORP8 contributes to phagosomal lipid regulation by Sec22b

Sec22 was reported to interact with LTPs Osh2 and Osh3 in yeast ^16^ which are homologs of the mammalian ORP family of LTPs ^35, 39^. In mammals, the closely related ORP5 and ORP8 isoforms mediate lipid exchange of PI(4)P at the PM as well as at endo-lysosomal compartments ^35, 40^. Thus, we asked whether ORP5 or 8 could contribute to Sec22b-regulated lipid exchange at phagosomes. Although several attempts to immunoprecipitate endogenous or overexpressed Sec22b were made in phagocytosing MEFs, the results were inconsistent. On the other hand, in HeLa cells where transfection efficiency was higher and more evenly distributed, FLAG-Sec22b was co-expressed with EGFP-tagged ORP5 or ORP8 or with cytosolic EGFP as control (Fig. 6A). Sec22b co-immunoprecipitated with both ORP5 and ORP8, but not with cytosolic EGFP, suggesting that interactions between Sec22b and LTPs are conserved in mammalian cells. In addition, phagocytic MEF cells were co-transfected with either EGFP-ORP5 or EGFP-ORP8 and mCherry-Sec22b. While Sec22b showed a typical reticular pattern, ORP5 was mostly found in peripheral patches reminiscent of ER-PM junctions ^40^ resulting in only minor overlap with Sec22b (Fig. 6B). In contrast, co-expression of mCherry-Sec22b and EGFP-ORP8 showed a striking overlap both in the ER and around phagosomes (Fig. 6B). We next investigated if ORP5/8 are involved in transporting lipids across ER-Phg MCS using siRNA. Although the siRNA pools employed were inefficient at downregulating ORP5 and 8 protein expression individually, when applied simultaneously a downregulation of 59% and 61% for ORP5 and 8 respectively was observed (Fig. S3B-D). The phagosomal PI(4)P dynamics were then assessed in ORP5/8 depleted cells as described above. Interestingly, ORP5/8 depletion resulted in a slightly but still significantly increased phagosomal PI(4)P levels, phenocopying the knockdown of Sec22b (Fig. 6C) which argues for a role of these LTPs in transporting PI(4)P from the phagosome to the ER. Moreover, overexpression of EGFP-ORP8, but not EGFP-ORP5 in Sec22b-depleted cells rescued the phenotype and reduced phagosomal PI(4)P to wild-type levels, similar to MAPPER overexpression (Fig. 6D). Since ORP8 overexpression is itself expected to increase MCS, to test whether the lipid transfer function was specifically required for the effect of ORP8 on phagosomal PI(4)P, a mutant EGFP-ORP8 harbouring two point mutations H514A-H515A (ORP8-Mut), which abrogates lipid transfer but not tethering ^40^ was employed. EGFP-ORP8-Mut showed a similar reticular/MCS expression pattern, though the mutant appeared to be more robustly recruited to PM-MCS and less reticular than the wild-type protein, and recruitment to phagosomes was less prominent (Fig. 6E). Although both ORP8 and ORP8-Mut reduced the initial PI(4)P peak, shSec22b-MEFs expressing the mutant ORP8 displayed significantly higher levels of phagosomal PI(4)P than wild-type ORP8, indicating the lipid transfer function of ORP8 is required for its ability to localize to phagosomes and to rescue the Sec22b knockdown phenotype. These data indicate that loss of Sec22b can be compensated by overexpressing ORP8, and that ORP8 does not require Sec22b to be recruited to MCS, although we cannot exclude that such a recruitment maybe more efficient in the presence of Sec22b. Taken together, our data point towards a function of Sec22b in the formation or stabilization of ER-Phg MCS and the recruitment of ORP8 which transports PI(4)P from the phagosome to the ER and PS from the ER to phagosomes representing a new function of ER-Phg MCS.

**Figure 6.**
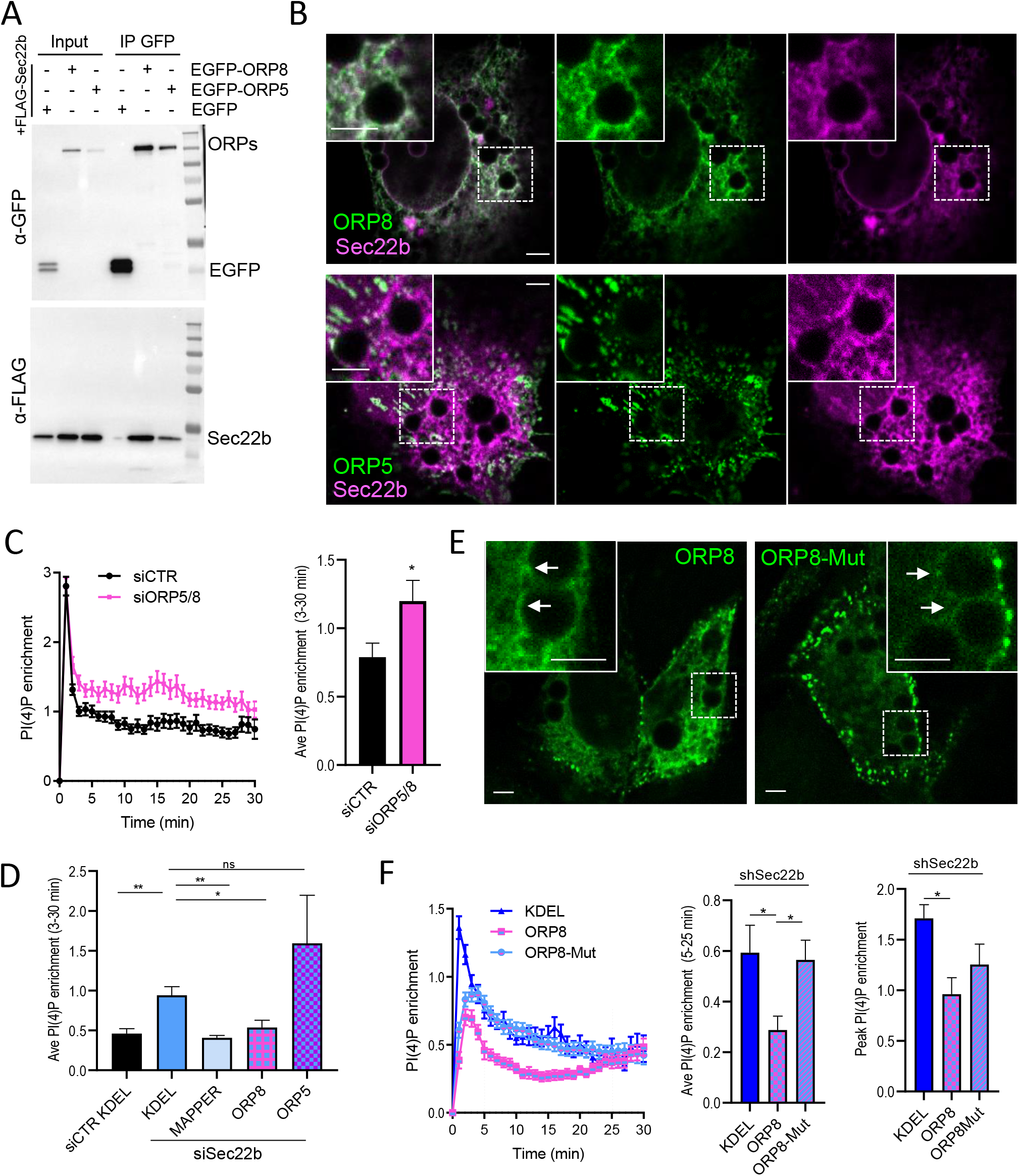
Sec22b interaction with ORP8 but not ORP5 regulates phagosomal PI(4)P. **A)** Western blot analysis of whole cell lysates (input) and GFP-trap agarose beads immunoprecipitates (IP) of HeLa cells transfected with FLAG-Sec22b and either EGFP-ORP8, EGFP-ORP5 or EGFP control, using anti-GFP and anti-FLAG. **B)** Confocal images of phagocytic MEFs co-transfected with mCh-Sec22b (magenta) and either EGFP-ORP8 (green, top panels) or EGFP-ORP5 (green, bottom panels), after 30 min exposure to IgG-beads. **C)** Increased levels of phagosomal PI(4)P between 3-30 min after target ingestion in WT MEFs transfected with GFP-2xP4M and either siCTR (100 nM) or siORP5/8 (50 nM siORP5 + 50 nM siORP8; n=9/11 siCTR/siORP5/8; see also Fig. S3B-D). **D)** Quantification of phagosomal PI(4)P between 3-30 min after IgG-bead ingestion in WT phagocytic MEFs transfected with mCh-2xP4M and siCTR+GFP-KDEL and siSec22b+GFP-KDEL, +GFP-MAPPER, +EGFP-ORP8 or +EGFP-ORP5. Phagosomal PI(4)P levels were higher in siSec22b+KDEL compared to siCTR+KDEL. In siSec22b-transfected cells MAPPER and ORP8 but not ORP5 decreased phagosomal PI(4)P as compared to KDEL controls. (n=10, 13/4/6/4 coverslips siCTR+KDEL, siSec22b+KDEL/MAPPER/ORP8/ORP5). **E)** Confocal images of shSec22b phagocytic MEFs transfected with EGFP-ORP8 or EGFP-ORP8-H514A-H515A (ORP8-Mut) after 30 min exposure to IgG-beads. **F)** Quantification of phagosomal PI(4)P peak levels and between 5-25 min after IgG-bead ingestion in shSec22b phagocytic MEFs transfected with mCh-2xP4M and GFP-KDEL, EGFP-ORP8 and EGFP-ORP8-Mut. ORP8 but not ORP8-Mut significantly decreased phagosomal PI(4)P as compared to KDEL controls (n=7/11/10 KDEL/ORP8/ORP8-Mut). White bars = 3 µm. Bar graphs are means +SEM. * p<0.05, **p<0.01.

### Sec22b-mediated tethering controls phago-lysosome fusion

We next investigated the physiological relevance of this lipid transfer mechanism at ER-Phg MCS. PI(4)P has been shown to promote phagosome maturation and phago-lysosome fusion by recruiting Rab7 effectors ^32, 41^. As discussed above, increased phagosomal LAMP1, and an exacerbated degradation of ingested antigens was observed upon Sec22b knockdown in DCs, suggesting an increase in phago-lysosome fusion, although no mechanism was provided ^19^. We thus employed an established Förster resonant energy transfer (FRET)-based phago-lysosome fusion assay ^7, 42^ to examine this process in our model system. Phagocytic MEFs were loaded with the impermeant FRET acceptor Alexa Fluor (AF) 594-HA which accumulated in lysosomes. The cells were then exposed to IgG beads coupled to the FRET donor AF488, and the FRET, total acceptor and total donor signals were measured by microscopy. In congruence with previous reports, after 90 min of phagocytosis, phago-lysosome fusion (PLF index) was significantly increased in Sec22b-depleted cells as compared to controls (Fig. 7A), whereas the rate of phagocytosis (phagocytic index) and total lysosomal loading was similar between the two conditions (Fig. 7B). Phagosomal pH was also similar upon Sec22b knockdown (Fig. 7C), which is consistent with results observed in DCs ^19^, with previous reports suggesting that V-ATPase delivery to phagosomes precedes phago-lysosome fusion ^43, 44^, as well as our own observations that when phago-lysosome fusion is decreased in *Stim1^-/-^* cells, pH remained unchanged ^7^. We then sought to determine whether the PLF index could be rescued by overexpression of Sec22b or different MCS tethering and control constructs to verify whether the effects observed on PI(4)P mirrored effects on phago-lysosome fusion. Surprisingly, the first observation was that transfection/overexpression of KDEL alone appeared to reduce the PLF index in general (compare Fig. 7A with Fig. 7D) with perhaps a greater effect in shSec22b cells, leading to a diminished dynamic range, and loss of a significant difference between the shCTR+KDEL and shSec22b+KDEL conditions (Fig. 7D). On the other hand, overexpression of shR-Sec22b significantly lowered the PLF index compared to KDEL in shSec22b cells at both low and high phagocytic rates (Fig. 7D-F and Fig. S4A-B respectively). Although GFP signals represented only at most 10% of the much brighter donor dye (AF488) channel signal and were mostly excluded by a restrictive phagosomal segmentation, one caveat of this technique is that differences in construct expression levels cannot be entirely excluded from the PLF index calculation. While most constructs appeared to be expressed at similar levels, expression levels of GFP-KDEL were visibly lower than EGFP-Sec22b, and thus we also calculated a FRET ratio independently of the green channel fluorescence (FRET signal normalized to total acceptor dye loading). In this case we still observed a significant decrease upon expression of Sec22b (Fig. S4A-B), supporting the idea that Sec22b overexpression indeed directly lowers phago-lysosome fusion. In contrast, although a trend towards a partial effect could be discerned, the PLF index upon overexpression of shR-Sec22b-P33, in the context of similar phagocytic index and total acceptor loading, was not significantly different from that of KDEL, indicating that MCS gap disruption at least in part contributes to this function (Fig. 7C-E). Moreover, both MAPPER and ORP8 but not ORP8-Mut significantly lowered the PLF index without affecting phagocytic rates, indicating that lowering phagosomal levels of PI(4)P through MCS tethering also impacts phago-lysosome fusion (Fig. 7F-G). However, in the case of MAPPER and ORP8-Mut the total lysosomal loading of acceptor dye was slightly higher than with KDEL, suggesting more generalized effects on the endocytic pathway may contribute to this phenotype. However, since MAPPER and ORP8-Mut had opposite effects on the PLF index while having similar effects on total acceptor loading, differences in the latter cannot fully explain the changes in PLF observed. Thus, while other mechanism likely contribute, together these data suggest that Sec22b regulates phagosome maturation at least in part by recruiting ORP8 to phagosomal MCS and mitigating levels of PI(4)P.

**Figure 7.**
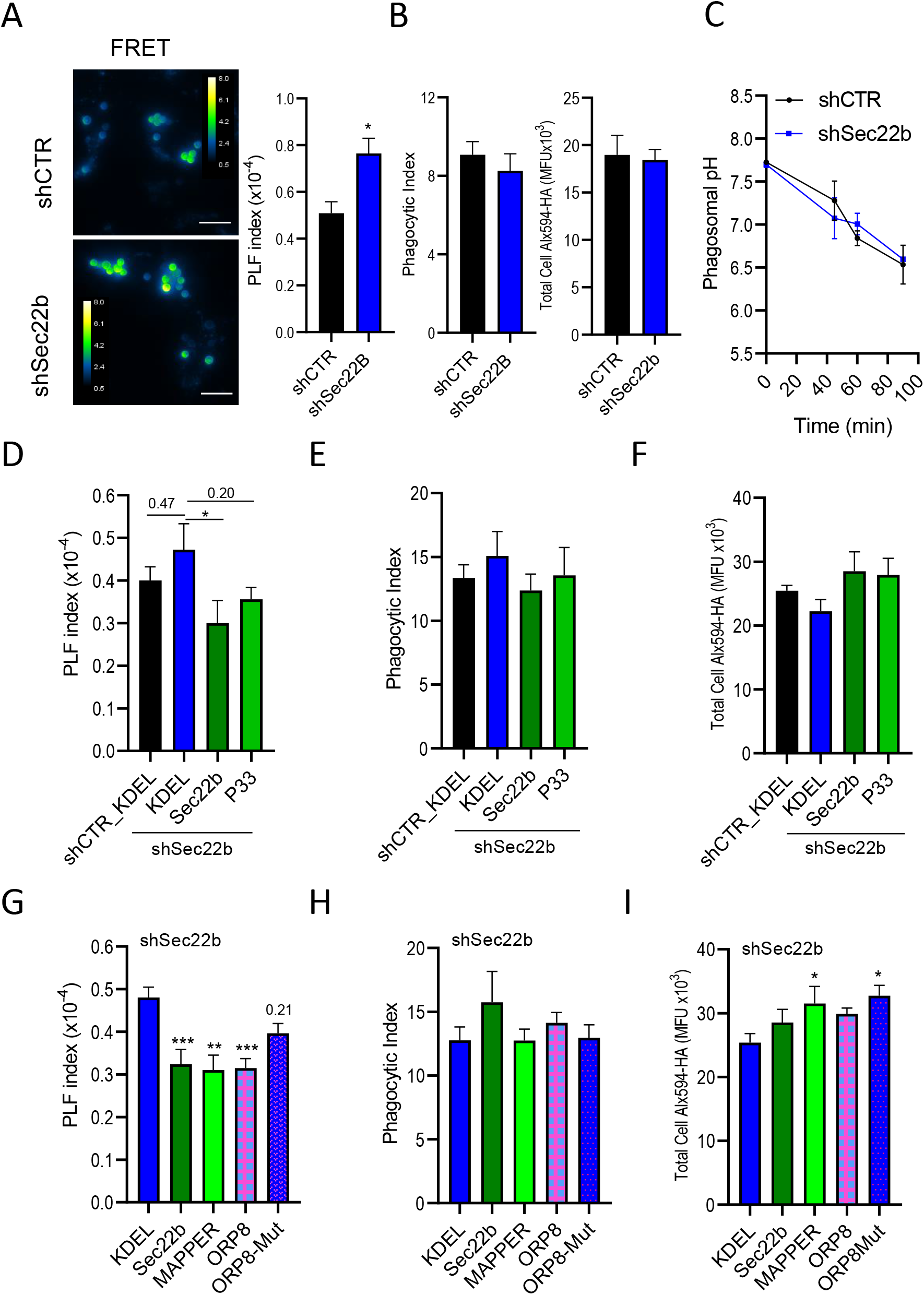
Sec22b MCS tethering regulates phago-lysosome fusion. **A-B)** Phago-lysosome fusion (PLF index, see methods) measured in phagocytic shCTR and shSec22b MEFs pulse-chased with 10 µg/ml FRET acceptor AF594-HA and exposed to FRET donor-coupled AF488-IgG-beads for 90 min. PLF index is increased in shSec22b cells (right panel, n= 6). The left panels show representative images of the maximum projection of FRET signal stacks (480/630 ex/em, 15 planes/0.8 µm spacing) of shCTR (top panel) and shSec22b (bottom panel) in ratiometric green-blue pseudocolor at identical contrast settings. **B)** Quantification of the number of phagosomes per cell (phagocytic index, left panel) and total cell loading of AF549-HA (right panel) of the cells analysed in (A). **C)** Phagosomal pH was measured by ratiometric (440/480) imaging in shCTR and shSec22b phagocyotic MEFs after 40, 60, and 90 min exposure of cells to IgG-FITC-RBC. **D)** PLF index measured in phagocytic shCTR and shSec22b MEFs transfected with GFP-KDEL, shR-EGFP-Sec22b or shR-EGFP-Sec22b-P33 as indicated. In shSec22b cells overexpression of Sec22b reduced PLF index as compared to KDEL controls (n=10;5/6/6 for shCTR+KDEL;shSec22b+KDEL/+Sec22b/+P33). **E-F)** Quantification of phagocytic index (E) and total cell loading of AF549-HA (F) of the cells analysed in (D). (See also Fig. S4A-B). **G)** PLF index measured in phagocytic shSec22b MEFs transfected with GFP-KDEL, shR-EGFP-Sec22b, GFP-MAPPER, EGFP-ORP8 or EGFP-ORP8-Mut. In shSec22b cells overexpression of Sec22b, MAPPER and ORP8 reduced PLF index as compared to KDEL controls (n=11/6/15/7/7 KDEL/Sec22b/MAPPER/ORP8/ORP8-Mut). **H-I)** Quantification of phagocytic index (E) and total cell loading of AF549-HA (F) of the cells analysed in (G). Total cell loading was increased in MAPPER and ORP8-Mut-transfected cells as compared to KDEL controls. White bar = 10 µm. Bar graphs are means + SEM. *p<0.05, **p<0.01, ***p<0.001.

## Discussion

When we first observed STIM1-dependent ER-phagosome MCS in neutrophils we found a strong association between these structures and localized Ca^2+^ hotspots ^5^. Yet residual hotspots persisted in cells lacking STIM proteins, and at least 50% of the signal originated from Ca^2+^ released from ER by IP_3_R, leading us to seek other MCS tethers ^6^. Although global Ca^2+^ signals in HeLa cells were unchanged upon Sec22b knockdown, expression of the Sec22b-P33 mutant reduced luminal ER Ca^2+^ and delayed Ca^2+^ refilling ^16^, suggesting that Sec22b may indirectly affect Ca^2+^ handling. We thus investigated whether a connection between Sec22b and STIM1 might exist either at the PM or phagosomes. In contrast to the observed effects in HeLa, in MEFs, Sec22b knockdown as well as P33 overexpression both increased global cytosolic Ca^2+^ signals (Fig. 3F, G), whereas overexpression of Sec22b had a dramatic effect on STIM1 recruitment that surprisingly did not translate to changes in cytosolic signals. Conceivably, Sec22b stabilizes MCS allowing a faster infiltration of STIM molecules, yet its presence may interfere with STIM-ORAI pairing either by recruiting components that displace these Ca^2+^ channels, by changing the local lipid environment, or perhaps by regulating MCS gap distance, parameters shown to modulate Ca^2+^ channel recruitment and activity ^26, 45, 46^. Such parameters may also be cell type-dependent, explaining differences between this and previous reports. In contrast to the effects on global signals, the frequency of local signals was not affected by Sec22b downregulation (Fig. 4). However, since the low levels of BAPTA required to visualize local hotspots precludes accurate estimation of the peak Ca^2+^ concentration within hotspots, a more subtle effect on hotspot Ca^2+^ content may have remained undetected. Thus, we cannot completely exclude that an increase in the size (rather than frequency) of Ca^2+^ hotspots could contribute to increased phago-lysosome fusion upon Sec22b depletion, though the dramatic effect on PI(4)P (Fig. 5) seems a more likely explanation. On the other hand, the reported inhibition of ER Ca^2+^ refilling upon Sec22b-P33 expression ^16^ could explain why Sec22b-P33 overexpression still shows a tendency towards reduced phago-lysosome fusion (Fig. 7C), despite the complete inability to rescue levels of PI(4)P in Sec22b-depleted cells (Fig. 5F). Regardless, it should be kept in mind that Sec22b may have subtle effects on Ca^2+^ signaling which may be more or less important depending on cell type.

In DCs, Sec22b knockdown induced a clear increase in LAMP1 recruitment and in phagosomal degradation ^19^. In our hands increased phago-lysosome fusion was not due to off-target effects, as has been suggested for the shSec22b effect on cross-presentation ^47^, since different sh- or siRNAs produced the same phenotype, and Sec22b re-expression significantly lowered levels of phago-lysosome fusion in knockdown cells (Fig. 7). Our initial hypothesis was that this occurs by a Ca^2+^-related mechanism, since Ca^2+^ is well-known to promote phagosomal maturation through the activation of Ca^2+^-dependent proteins such as synaptotagmins, calcineurin and annexins among others ^48^. However, our data suggest the mechanism through which this occurs involves instead the regulation of phagosomal phospholipids through the recruitment of LTPs at phagosomal MCS. We found that genetic manipulations that changed levels of phagosomal PI(4)P in shSec22b cells, such as expression of MAPPER and ORP8 which have been reported to only minorly impact cytosolic Ca^2+^ ^26, 49^, correlated well with changes in phago-lysosome fusion. This is in line with previous observations that this lipid serves as a docking site for Rab7 effectors such as RILP thus promoting lysosomal docking and fusion ^32, 41^. However, we also detected decreases in phagosomal PS and PI(3)P, which may contribute to changes in phagosomal maturation. Whereas phagosomal PS has been linked to surface charge and recruitment of c-Src, PI(3)P recruits effectors such as VPS34 and PI(3,5)P2-kinase PikFYVE promoting phago-lysosome fusion ^38, 50, 51^. Their decrease would thus be predicted to reduce rather than increase fusion with endocytic vesicles. In contrast, PI(3)P binding to the NADPH oxidase subunit p40*^phox^*(NCF4) promotes phagosomal reactive oxygen species (ROS) production, which could serve to delay phagosomal maturation by preventing acidification, and is critical for efficient cross-presentation ^44, 52, 53^. However, in MEFs ROS production is presumably negligible, and in macrophages Sec22b knockdown did not impact Nox2-mediated ROS production ^54^, although it did reduce iNOS-^55^ and IRE1α-^54^ dependent ROS at late (<8 hr) timepoints in DCs and macrophages respectively. How Sec22b regulates PI(3)P is still obscure, although a feedback coordination with PI(4)P could be involved. Whether changes in PI(3)P also occur and could contribute to the knockdown phenotypes in DCs would certainly be an interesting question to pursue in future studies. In addition, whether other phagosomal lipids such as cholesterol or ceramides, which also depend on phagosomal MCS ^56, 57^, are impacted by Sec22b, and whether this is similar in different phagocytic cell types would also be interesting questions to address.

Finally, the two critical remaining questions are what is the impact of Sec22b-regulated phago-lysosome fusion on cross-presentation, and how does this relate to ERGIC fusion? Yet answering these questions is not trivial. Firstly, although reduced phagosomal proteolysis has been suggested to favor cross-presentation by limiting the complete destruction of antigenic peptides ^52, 53, 58^, clearly some processing is still required since blocking phagosomal acidification and protease activity also negatively impact cross-presentation ^58–60^. Indeed, we and others have correlated the increased delivery of the endosomal protease IRAP to improved cross-presentation ^7, 61^, highlighting the importance of a careful balance. Model phagocytes similar to our MEFs are capable of cross-presenting antigens even if less efficiently than DCs ^62^. In principle, they can offer advantages beyond easier and cheaper genetic tractability, since factors such as cytokine secretion or cell-surface expression of co-stimulatory or co-inhibitory molecules, which all contribute to T cell responses are likely to play a lesser role in MEFs than DCs. Indeed IL-1β, IL-6 and TNFα secretion have been recently shown to be regulated by Sec22b ^55, 63^. However, our efforts in addressing this question in MEFs have been complicated by an interesting and surprising effect of Sec22b downregulation, which increases by nearly 4-fold the steady-state levels of cell surface MHC-I (Fig. S4C-D). This phenotype is at least in part linked to Sec22b function as re-expression of Sec22b partially reversed the effect (Fig. S4E-F). We are currently investigating the mechanisms underlying this effect, as well as whether this mechanism is also operational in DCs where MHC-I trafficking may be different ^64^.

Nevertheless, the impact of Sec22b on cross-presentation, which has been recently contested ^47^, has up to now always been assumed to rely on its ability to deliver ER proteins to phagosomes through fusion of ERGIC vesicles ^19, 64–66^. Here, we provide evidence that both endogenous and overexpressed Sec22b tether and regulate MCS formed between the ER and phagosomes, where the use of 3D-CLEM confirms a very close but non-fusogenic interaction of Sec22b containing membranes. MCS are non-fusogenic, stable associations between organelles that survive biochemical fractionation, and protocols for the biochemical isolation of the most well-studied MCS, that of ER-mitochondria contacts, are well established^9, 67^. In our hands, fluorescent, STIM1-containing MCS were still visible in isolated phagosomes treated with ATP (shown to help shed contaminants ^68^) as well as 2 h of proteinase K digestion, suggesting MCS are difficult to remove (unpublished results). Our observations in this manuscript therefore now beg a re-interpretation of previous studies investigating the Sec22b-mediated recruitment of ER proteins to phagosomes, since at the very least a portion of the ER proteins detected are likely still contained within MCS rather than on the phagosomal membrane itself. Indeed, this might explain the punctate rather than continuous immunostaining of antigen-loaded MHC-I molecules around phagosomes ^69^, the presence of “empty” or endoglycosidase H-sensitive MHC-I molecules on isolated phagosomes ^66, 70^, and more recently the major loss of Sec22b on isolated phagosomes after 1 h proteinase K digestion ^55^. Our results do not rule out that Sec22b mediates fusion of ERGIC vesicles with phagosomes, and indeed we observe a close association between phagosomes and structures containing both Sec22b and ERGIC-53 (Fig. 1A, B, D) even in MEFs, as has been previously reported in DCs ^19, 55, 64^. We are currently developing more precise methods to measure ERGIC fusion in order to assess the relative contribution of Sec22b to this process in MEFs and DCs. As discussed previously ^4^, other evidence of direct communication between the ER (or ERGIC) also support the direct delivery of ER proteins to phagosomes. Nevertheless, a change in Sec22b-mediated MCS formation may confound measurements of factors presumed to increase or decrease ERGIC fusion with phagosomes. Sec22b tethering, involving SNARE complexes with Stx1, 3 or 4 in the absence of SNAP-25, versus its fusogenic propensity may in fact be regulated by its N-terminal longin domain ^15^, or potentially by post-translational modifications such as ubiquitination ^71^. Thus, a shift in the balance of MCS versus fusion may help explain discrepancies between prior studies, and future studies that distinguish between effects on MCS and fusogenic pathways may help formulate a more accurate picture of the complex biology of this important trafficking regulator.

## Materials and Methods

### Reagents

The following antibodies (antibody name/catalog#/dilution) were purchased from: Synaptic Systems (SYSY): rabbit anti-Sec22b (186003/1:1000); Santa Cruz: mouse anti-Sec22b (29-F7) (sc-101267/1:100); Cell Signaling: mouse anti-c-myc antibody (9B11) (2276/1:100), rabbit anti-STIM2 (4917S/1:1000); Thermo: mouse anti-CD16-CD32 (Fc-Block, 14-0161-85, 1:200), rabbit anti-ORP5 (PA5-18221/1:500), mouse-anti-MHC Class I-PE (H-2Kb, AF6-88.5.5.3, 12-5958-80, 1:100) goat-anti-rabbit Alexa Fluor 555 (A21428/1:1000), goat-anti-mouse Alexa Fluor 647 (A21235/1:1000); GeneTex: rabbit anti-ORP8 (GTX121273/1:500); Sigma: mouse anti-α-tubulin (T9026/1:5000), rabbit anti-sheep red blood cell (sRBC, S1389/1:40), mouse anti-FLAG-M2 (F1804/1:1000); mouse anti-GFP (11814460001/1:1000); BD Biosciences: mouse anti-GOK/STIM1 (610954/1:100); Bio-Rad: goat anti-rabbit IgG (H+L) HRP conjugate (170-6515/1:10000), goat anti-mouse IgG (H+L) HRP conjugate (170-6516/1:10000), Innovative Research: human IgG protein A purified (hIgG) (IR-HU-GF); Jackson ImmunoResearch: donkey-anti-mouse Alexa Fluor 488 (715-545-150/1:800), goat-anti-rabbit Alexa Fluor 647 (111-605-003/1:1000). Interfering RNA was purchased from: Santa Cruz: mouse siSec22b (sc-153306, siRNA pool/50 nM), siCTR (sc-37007/50-100nM); Dharmacon/GE Healthcare: ON-TARGETplus siRNA mouse siORP8 (Osbpl8 (237542) SMARTpool SO-2755574G/50 nM), ON-TARGETplus siRNA mouse siORP5 (Osbpl5 (79169), SMARTpool so-2791415G/50 nM); mouse shSec22b lentiviral clone (TRC Clone ID: TRCN0000115089); Sigma: shCTR non-target control particles SCH002V Mission shRNA Lentiviral clone. All lentiviral particles were produced in Lenti-X 293T cells using the Lenti-X HTS Packaging System (Takara, Japan) according to the manufacturer’s instructions. Lentiviral titers were determined using the LentiX-p24 Rapid Titer ELISA kit (Takara). All cell culture reagents were obtained from ThermoFisher Scientific, and all chemicals were purchased from Sigma-Aldrich unless otherwise stated.

### Plasmids

The following plasmids were a gift from the laboratory of Prof. Nicolas Demaurex (University of Geneva): pcDNA3-myc-FCGR2A (Fc receptor) ^50^, pEGFP-FCGR2A ^50^ p-DS-XB-GFP-MAPPER-Long ^26^, pCMV/Myc/ER-GFP (KDEL-GFP) (Thermo), pAmaxa-ER-TagRFP (RFP-KDEL) ^72^, YFP-STIM1 (Addgene plasmid # 18857)^73^, GFP-C1-PLCdelta-PH (Addgene plasmid # 21179)^34^. Plasmids pEGFP-ORP5, pEGFP-ORP8 and pFLAG-Sec22b were described previously ^74^. GFP-ERGIC-53 was a gift from Drs. Houchaima Ben Tekaya and Dr. Jean Gruenberg (University of Geneva) ^75^. pCMV-EGFP-Sec22b, pCMV-EGFP-Sec22b-P33, pCMV-mCherry-Sec22b were gifts from Drs. Thierry Galli and Christian Vannier (INSERM, Paris) ^16^. mRFP-Lact-C2 (Addgene plasmid # 74061)^38^, GFP-and mCherry-2xP4M ^31^ were kind gifts from Dr. Sergio Grinstein (University of Toronto). pRS424GFP-FYVE(EEA1) was a gift from Scott Emr (Addgene plasmid # 36096). TagRFP-FYVE(EE1A) was subcloned from pRS424GFP-FYVE(EEA1) into pTagRFP-C (Evrogen) and pEGFP-C1 (Clontech) using EcoRI/KpnI restriction enzymes (NEB). EGFP-Sec22b-shR and EGFP-Sec22b-P33-shR were generated by site-directed mutagenesis (F: CTTCTGAATGAAGGTGTCGAACTCGATAAAAGAATAAGGCCTAGACACAGTGGGC; R: GCCCACTGTGTCTAGGCCTTATTCTTTTATCGAGTTCGACACCTTCATTCAGAAG) using the Q5 Site-Directed Mutagenesis Kit (NEB). pEGFP-ORP8-H514A-H515A (ORP8-Mut) was similarly generated (F: ACAGGTGTCCgctgctCCACCAATATCTG, R: TCAGCAATATAAAAAGTTTTGC).

### Cell culture, transfection and transduction

Wild-type mouse embryonic fibroblasts (MEF, ATCC CRL-2991), *Stim1^−/−^* (SKO) MEFs ^76^, *Stim1^−/−^;Stim2^-/-^* (DKO) MEFs ^77^, and HeLa cells were grown in DMEM (22320) containing 10% fetal calf serum, 0.5% penicillin/streptomycin (pen/strep), at 37°C and 5% CO_2_ and were passaged twice a week. MEF cells were used between passages 5 and 50. Cells were transfected using Lipofectamine 2000 in full medium without antibiotics for 4-6 h with cells at 50-60% 1-2 days after seeding for plasmids or at the same time as seeding (reverse transfection) for transfections containing siRNA. Lentivirus transduction was performed by centrifuging cells and viral particles at 5 MOI in complete medium supplemented with 8 μg/mL polybrene at 500 g at 37 °C for 1 h. To obtain stable cell lines, puromycin (10 µg/mL) selection was performed 2 days after transduction. Stable cell lines (shCTR, shSec22b) were maintained in medium supplemented with (10 µg/mL) puromycin.

### Phagocytic target preparation

3 μm carboxyl polystyrene microspheres (Spherotech/ Cat No. CP-30-10) were covalently coupled with hIgG by washing 3 times in sterile PBS at maximum speed (18,000 g) at 4°C, activating with 50 mM 1-Ethyl-3-(3-dimethylaminopropyl)carbodiimide hydrochloride(EDC-HCl) (Carl Roth/2156.1) for 15 min in PBS at room temperature (RT) with rigorous shaking, followed by 3 washes at 4°C in 0.1 M Na_2_B_4_O_7_ buffer (pH 8.0). 6 mg of hIgG were added to the beads and incubated overnight at 4°C on a shaker. The following day, the beads were washed 2x with 250 mM glycine/PBS followed by two washes in PBS. For fluorescent IgG-bead preparation 20 μg/mL Alpha Fluor 488 amine (AF488-Amine) (AAT Bioquest/Cat No. 1705) were added after EDC activation, and particles were incubated with the dye for 30 min at RT with agitation followed by one wash before adding hIgG. Gluteraldehyde-stabilized sheep RBCs were incubated for 1h in 0.1% NaBH_4_, washed 3x in 0.1M Na_2_CO_3_ (pH 9.3 or pH 8.0) followed by incubation with 1 mg/ml fluorescein isothiocyanate (FITC) or pHrodo Red, succinimidyl ester (Thermo/P36600), respectively, for 4 h and washed 3x in PBS. Sheep RBCs were opsonized in rabbit anti-sRBC for 1h at 37°C, followed by 3 washes in PBS at 4°C just prior to use. Prepared beads and RBCs were resuspended in PBS+0.5%pen/strep, counted using a Countess (Thermo) cell counter, and added to cells at 10:1 target:cell ratio unless otherwise indicated. All buffers were sterile filtered using 0.2 µm filters.

### Immunofluorescence

Cells seeded on 0.17 thickness 12 mm coverslips (Carl Roth) were fixed in 4% paraformaldehyde (PFA, Electron Microscopy Sciences)/PBS for 30 min. For immunostainings, cells were permeabilized in 0.1% TritonX-100/PBS, reduced with fresh 0.1% NaBH_4_ (Carl Roth)/PBS for 10 min, treated with Image-IT-Fx (Thermo)for 30 min, blocked in 1%BSA/PBS (blocking buffer) for 30 min, and incubated overnight at 4°C in primary antibody and for 1 h at RT in secondary antibody. Coverslips were mounted in SlowFade Gold (Thermo). Images were acquired either using a Nikon A1r confocal microscope system/60x 1.4 CFI Plan Apochromat objective or a Zeiss LSM700 system/Plan-Apochromat 63x /1.4 objective. Z-stacks were taken at 0.5 µm intervals. Quantification of MAPPER recruitment to phagosomes was performed manually using ImageJ. A 0.2 µm (approx. 4 pixel) ring beyond the phagosomal border (based on the brightfield image), was examined for regions above threshold (manually defined per cell). The number of MAPPER punctate structures (> 0.01 µm^2^/4 pixel area) around each phagosome is reported.

### Electron Microscopy

Transmission electron microscopy (TEM) was performed at the Pole Facultaire de Microscopie Electronique at the University of Geneva as described ^5^ except uranyl acetate staining was replaced with Reynold’s (lead citrate, Electron Microscopy Sciences), and 20 nm sections were used. Grids were imaged on a Tecnai transmission electron microscope (FEI) and quantification was performed manually using ImageJ as described ^5^.

### Correlation Light Electron Microscopy (CLEM)

CLEM was performed as described ^7, 78^. Briefly, transfected cells seeded on Grid-500 polymer dishes (Ibidi) and exposed to phagocytic targets were fixed in 4% PFA for 30 min prior to high resolution confocal and brightfield imaging. After light image acquisition dish was re-fixed in 2.5% glutaraldehyde/2% PFA/2 mMCaCl_2_/0.15 M sodium cacodylate buffer (pH 7.4) for 3 h, dehydrated and embedded in Epon, and stained with osmium, ferrous cyanide, lead acetate and uranyl acetate as described ^7^. After sectioning the polymer grided coverslip was dissolved in xylol for 1 h, and samples mounted and sputter-coated with gold for 30 sec using a Q150T ES coater (Quorum Technologies). Focused ion beam scanning electron microscopy (FIB-SEM) imaging was performed on a Helios NanoLabG3 microscope (FEI). Images were acquired at the highest resolution setting, resulting in 5×5×10 nm pixels using the Autoslice and View software (FEI). Drift correction and alignment were performed using Amira software. Confocal stacks were deconvolved using Imaris (Oxford Instruments). Orthogonal slices and overlay were generated from the aligned stack using ImageJ.

### Calcium Imaging

Cells seeded on 25 mm coverslips were mounted on AttoFluor imaging chambers (Thermo), loaded with 4 μM Fura-2-AM, 0.01% pluronic (Thermo) in modified Ringer’s ^5^ for 30 min at RT. 340/380 nm excitation (ex) 510± 40 nm emission (em) ratiometric imaging was performed at 37 °C in modified Ringer’s where Ca^2+^-free solution (CF) contained 1 mM EGTA instead of 2 mM CaCl_2_ (CA). Images were acquired using a wide-field fluorescence Nikon Eclispe Ti microscope system (Visitron Systems) equipped with a 40X Plan Fluor 0.75 objective and stage heater. Frames were acquired every 3 s. Ca^2+^ microdomain/hotspot imaging was performed as described ^6^. Briefly, cells were loaded with 4 μM Fluo-8-AM (AAT Bioquest) for 30 min at 37 °C, 30 min at RT and 2.5 μM BAPTA-AM for the last 10 min, in modified Ringer’s/500 μM sulfinpyrazone. Simultaneous ex/em at 488/530 and 543/640 nm were collected in separate channels. Periphagosomal Ca^2+^ hotspots, images were averaged over 6 s and captured between 20 and 30 min after the addition of phagocytic targets. Image analysis was carried out with the aid of custom ImageJ macros on background subtracted images as described ^5, 6^.

### TIRF Imaging

Total internal reflection microscopy (TIRF) imaging was performed and quantified as described ^79^, except Cell Mask Deep Red (Thermo) was used to define the focal plane, experiments were carried out at 37 °C, and quantification was performed with ImageJ (NIH) and a custom macro. Briefly cells transfected and labelled with Cell Mask were rinsed 3x in 1 ml CF and imaged using a Nikon Eclipse Ti with a Perfect Focus system and 100x 1.49 Oil CFI Apochromat TIRF objective, at a rate of 1 frame/sec. MCS puncta were defined on background subtracted images as regions above threshold > 4 pixels.

### Phagosomal Phospholipid Assessment

Cells seeded on 25 mm coverslips and transfected with various lipid probes were mounted on AttoFluor chambers and washed in CA. Using a Nipkow Okagawa Nikon spinning disk confocal imaging system equipped with a temperature control chamber, stage motor and Plan Apo 63x/1.4 Oil DICIII objective and Visiview software (Visitron Systems). IgG-coupled phagocytic targets were added and 3-5 stage position were selected near cells that began phagocytosing with 5 min after bead addition. Confocal z-stacks of 9 frames spaced at 0.5 µm intervals were then acquired every 1 min at each stage position for 30-40 min. Images were taken in either 488/530, 561/630 nm ex/em or both depending on transfected probes. Background-subtracted maximum projections of confocal stacks were generated and initial total cell fluorescence as well as individual phagosome tracks analyzed using ImageJ custom macros. Phospholipid enrichment was defined as the phagosomal fluorescence divided the initial total cell fluorescence minus 1, and computed using Excel (Microsoft).

### Phagosomal pH

Phagosomal pH was assessed by ratiometric imaging as described ^80^, except opsonized fluorescein isothiocyanate (FITC)-coupled sRBCs were employed as phagocytic targets. Briefly, transfected cells seeded on 25 mm coverslips were mounted in AttoFluor chambers, washed in medium, exposed to phagocytic targets, centrifuged at 200 g and incubated at 37°C. After 15 min coverslips were washed 5x medium to remove unbound targets. At 35 min cells were washed in CA and 5 images at 440/530 and 480/530 nm ex/em were captured after 40, 60 and 90 min of target addition using the microscope as for Ca^2+^ imaging described above. Calibrations were performed using nigericin/monensin and potassium chloride buffers of pH 4-9 as described ^80^. Image analysis on background subtracted images was performed with ImageJ using custom macros, and pH calculations from calibration curves were computed using Excel.

### Phago-lysosome fusion (PLF) assay

Phago-lysosome fusion assays based on fluorescence resonance energy transfer (FRET) was performed as previously described ^7, 42^. Briefly, transfected cells were loaded with 10 µg/ml AF594-hydrazide (AF594-HA, Thermo) overnight and chased for 3 h into lysosomes. AF488-IgG beads were added, centrifuged, incubated for 30 min, and washed to remove un-internalized beads. Images were recorded using the microscope as for Ca2+ imaging described above except the 60x Plan-Apo 1.4 NA objective was used, and of 490/630 (FRET), 490/525 (green), 572/630 (red) nm ex/em, as well as brightfield channels were acquired. Stack of 15 planes spaced 0.8 µm in 5-10 fields per condition were recorded, and background-subtracted images were analysed using custom ImageJ macros. PLF index was defined per cell as the phagosomal fluorescence in the FRET/green channel ratio divided by the sum total cell acceptor loading as described ^7^.

### Immunoblotting

Cells were washed with cold PBS and resuspended in lysis buffer containing 25 mM Tris-HCl pH 7.6, 150 mM NaCl, 1% NP-40, 1% sodium deoxycholate, 0.1% SDS, in the presence of HaltProtease Inhibitor Cocktail, EDTA-Free (Thermo) and incubated on ice for 20 min. Lysates were sonicated for 10 min at 40 kHz in a bath sonicator (Emag Emmi-D280) and centrifuged at 18,000 g for 10 min at 4°C. Total protein concentration was determined using Roti-Quant kit (Carl Roth) according to the manufacturer’s instructions. Lysates were diluted in Roti-Load (Carl Roth) and heated at 95°C for 5 min. 15-35 µg protein were loaded and separated by SDS-PAGE in 4–20% Mini-Protean TGX Precast gels (BioRad). Gels were transferred to PVDF membranes using the iBlot system (Thermo). Membranes were blocked for 1 h in 5% low fat powdered milk/1% Tween/TBS (TBS-T), washed and incubated overnight in primary antibodies at 4°C and for 1h at RT in HRP-conjugated secondary antibodies all diluted in 3% milk/TBS-T. Signals were detected using Immobilon Western Chemiluminescent HRP Substrate (Millipore) and the ImageQuant LAS 4000 mini. Tubulin was used as an internal loading control, and the relative intensities of protein bands were quantified using ImageJ and Excel.

### Immunoprecipitation

Immunoprecipitation was performed using Chromotek GFP-trap agarose beads (Allele Biotech) as described ^74^. Briefly transfected cells were washed in cold PBS and lysed with buffer 50 mM Tris, 120 mM NaCl, 40 mM Hepes, 0.5% digitonin, 0.5% CHAPS, (pH 7.36) supplemented with a protease inhibitor cocktail (Roche). The cell lysates were then incubated with the beads for 1 h at 4^0^C in rotation. After extensive washes, the immunoprecipitated proteins bound to the beads were eluted in 2% SDS sample buffer and boiled for 1 min prior to SDS–PAGE in a 10% gel and immunoblotting as above.

### Flow Cytometry

Flow cytometry was performed as described ^7^. Briefly, cells were washed 1x in cold PBS detached for 15 min on ice in PBS/2 mM EDTA, centrifuged at 400 g, 5 min, and resuspended in FACS Buffer (2% BSA, 20 mM EDTA in PBS) + Fc-Block antibody for 30 min on ice. Cells were stained with anti-MHC-I-PE for 30 min, centrifuged and resuspended in FACS buffer. Analysis was performed using an Accuri C6 flow cytometer and CFlow Plus software (BD Biosciences).

### Statistics

Prism (GraphPad Software) was used to conduct all statistical analyses. An unpaired t-test with or without Welch’s correction was used to determine the statistical significance between two different groups depending on F test for variance similarity. A one-way ANOVA or Brown-Forsythe and Welch test with Dunnett’s post-hoc test was used to examine group sets, depending on F test for differences in variance. P values are labelled with asterisks: *P ≤ 0.05, **P ≤ 0.01, ***P ≤ 0.001 or listed above the bars.

## Author Contributions

NCS designed and performed experiments, analysed data and wrote the manuscript. SB and FG designed and performed experiments and analysed data. PNH conceived the project, designed and performed experiments and wrote the manuscript. FM, FB, and CC performed experiments and analysed data. MA performed experiments.

## Acknowledgements

We are grateful to the bioimaging, flow cytometry and electron microscopy core facilities at the University of Geneva Medical Centre for their invaluable help; to Drs. Nicolas Demaurex, Maud Frieden, Walter Reith and Amado Carreras-Sureda for their valuable advice; and Drs. Maria Cruz Cobo and Vladimir Girik for reading and editing the manuscript. This work was funded by a Young Investigator Subsidy from The Sir Jules Thorn Overseas Charitable Trust (to PNH), a Novartis Foundation grant (17B078, to PNH), a Professor Dr. Max Cloëtta Foundation Medical Researcher Grant (to PNH), a Swiss National Science Foundation grant (310030_189094, to PNH), as well as the FSER (FRM n°206548) and the FVA (eOTP:669122 LS 212527) grants to FG.

## Data availability

Full datasets will be made available on the University of Geneva’s data repository upon publication of the manuscript. In the meantime, upon reasonable request data may be made available by contacting the corresponding author.

## Abbreviations

ER: Endoplasmic reticulum
MCS: membrane contact site
ORP8: oxysterol-binding protein related protein 8
PI(3)P: phosphatidylinositol-3-phosphate
PI(4)P: phosphatidylinositol-4-phosphate
PI(4,5)P2: phosphatidylinositol-4,5-bisphosphate
PS: phosphatidylserine
SNARE: soluble N-ethylmaleimide-sensitive factor attachment proteins receptor
STIM: Stromal interaction molecule

---

**Figure S1.**
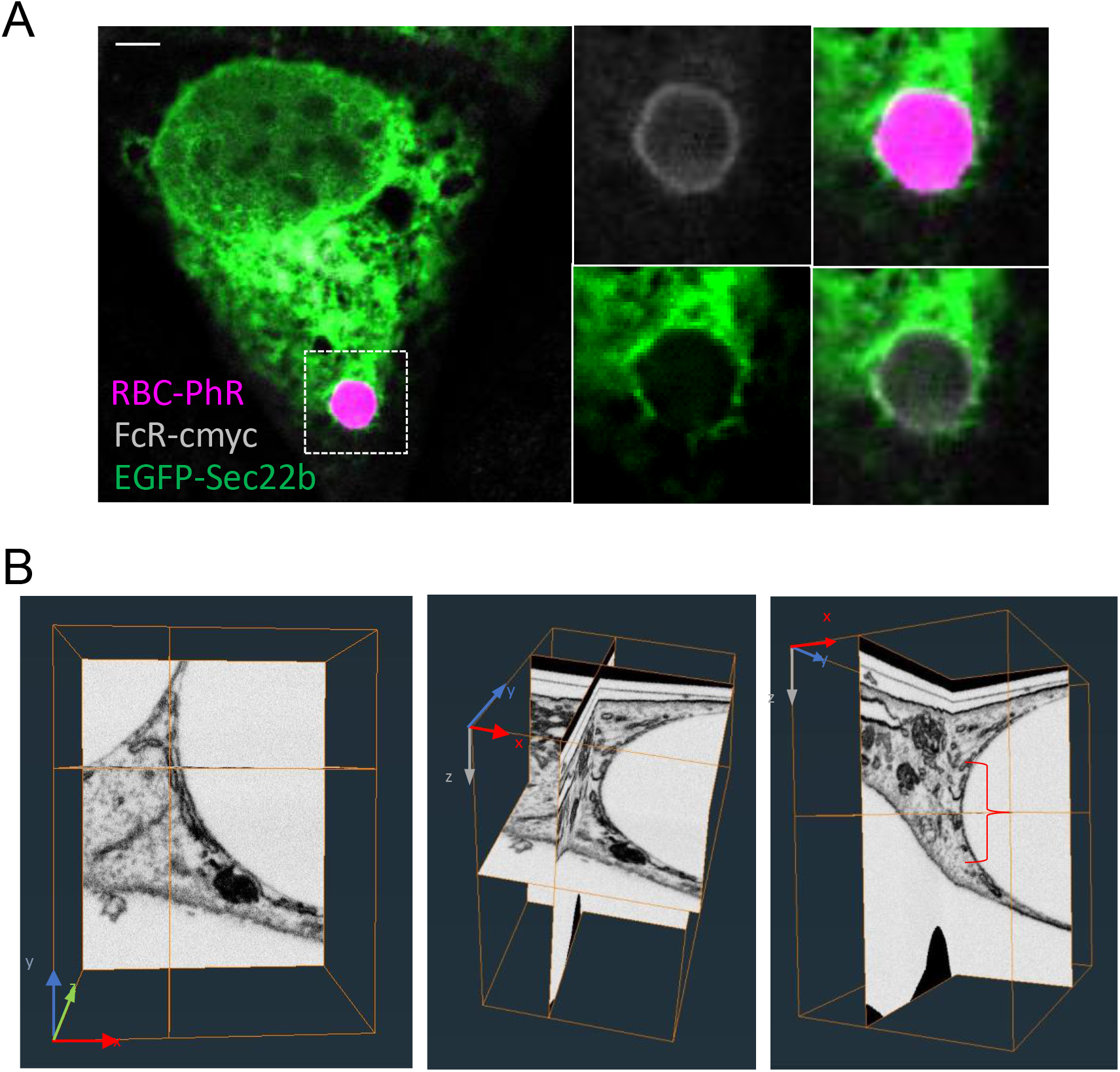
Related to Fig. 1. Sec22b localizes to ER-phagosome membrane contact sites. **A.** MEFs overexpressing FCGR2A(FcR)-c-myc (phagocytic MEFs, white) and EGFP-Sec22b (green) exposed to pHrodo-labelled IgG-sheep red blood cells (sRBCs, magenta) for 15 min and immunostained with anti-c-myc. **B)** The three panels show a sequential rotation (achieved by dragging the lower left-hand corner up) of orthogonal slices of the inset of the 3D EM stack shown in Fig. 1F. Examination of the area below and above (XZ) the plane of (XY) slice 261 shows that no vesicular structures that could correspond to ERGIC vesicles are present next to the MCS in the region spanning the fluorescent confocal slice (red bracket in right panel).

**Figure S2.**
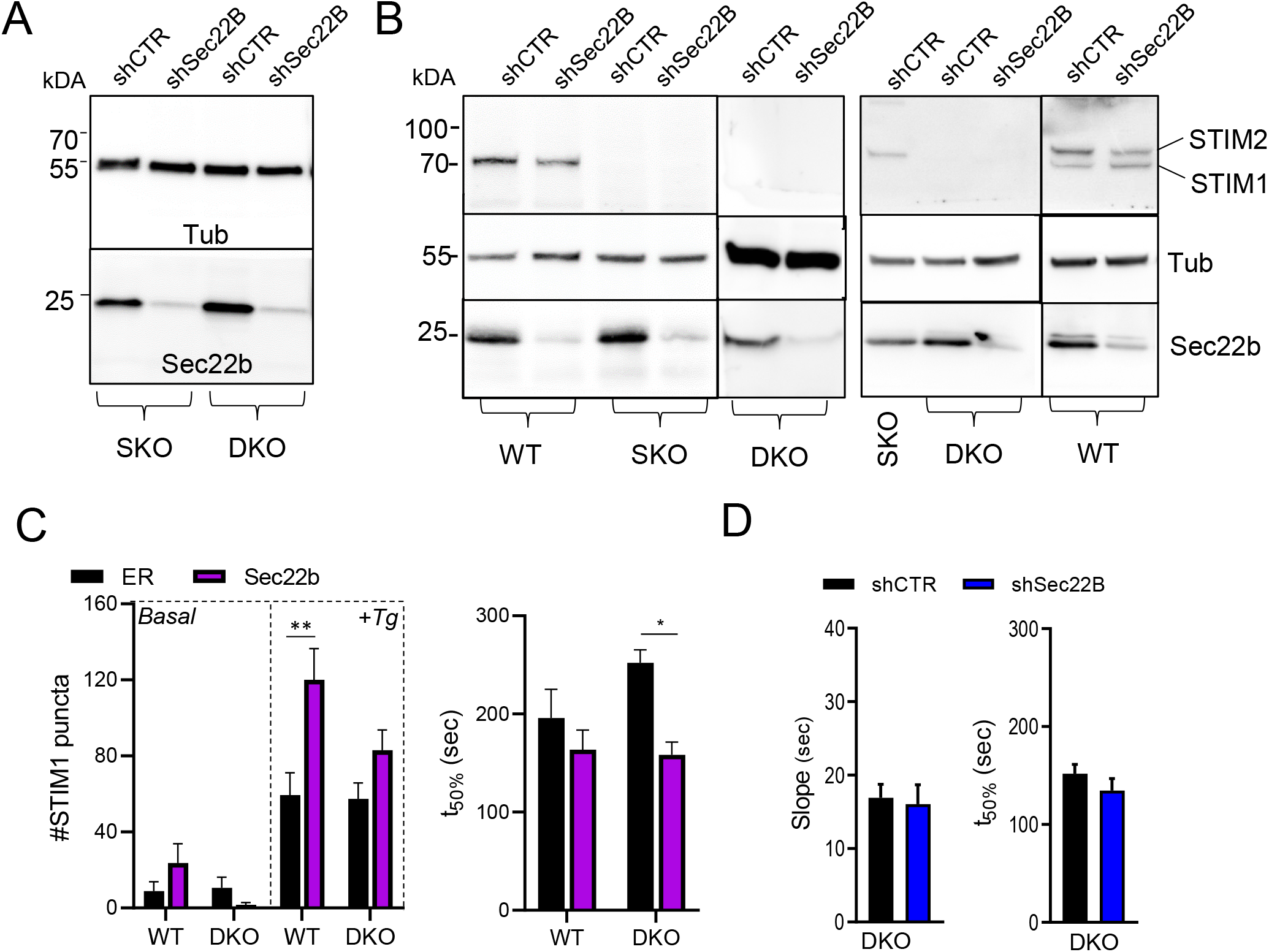
Related to Figs 2 and 3. Western blot and supplementary TIRF quantification. **A)** Representative Western blots of SKO (*Stim1^-/-^*) and DKO (*Stim1^-/-^; Stim2^-/-^*) MEF cell lines stably expressing shCTR and shSec22b incubated with anti-Sec22b (SYSY) and anti-tubulin as control show efficient Sec22b knockdown (quantification in Fig. 3I). **B)** Representative Western blots of WT, SKO and DKO stable cell lines incubated as above and with anti-STIM1 (left panel) or anti-STIM1 and STIM2 (right panel) confirm the genotype of the cells and show expression of STIM1 is not upregulated upon Sec22b knockdown (quantification in Fig. 3H). **C)** Related to Fig. 3A-C, quantification of the number (left panel) and kinetics (right panel, t_50%_ parameter of the Boltzman sigmoidal curve fit) of STIM1 puncta appearing at the TIRF plane after addition of Tg. In WT cells overexpression of Sec22b increases the number of STIM1 puncta, whereas in DKO cells the number is similar but the kinetics are faster. **D)** Quantification of the kinetics (slope and t_50%_ parameters of Boltzmann sigmoidal curve fits) of STIM1 puncta arrival at the TIRF plane in response to Tg shows Sec22b knockdown does not impact STIM1 recruitment. Related to Fig. 3D. Bar graphs show means +SEM, * p<0.05, **p<0.01.

**Figure S3.**
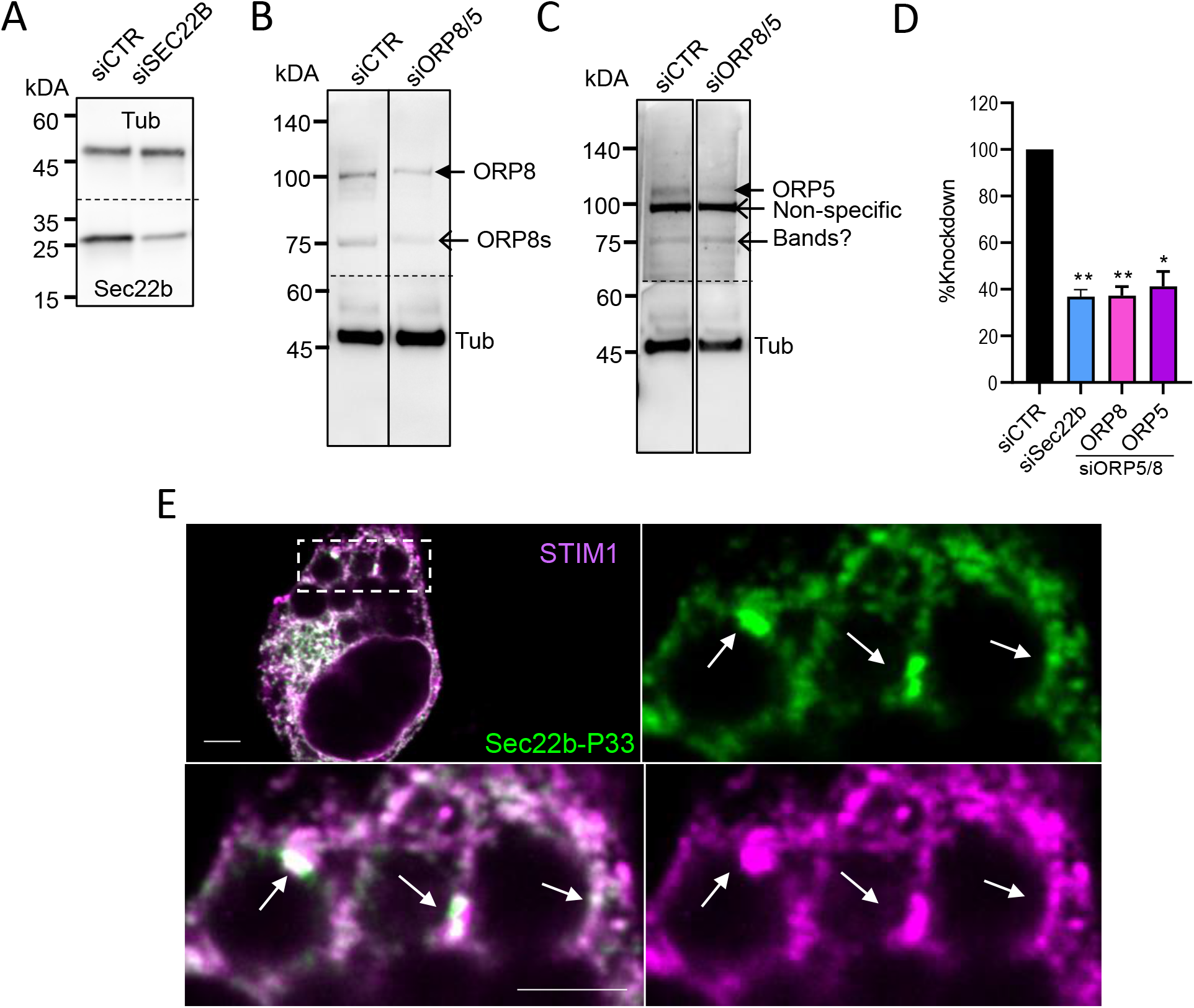
Related to Figs 5 and 6. Western blots for siRNA downregulation and Sec22b-P33 recruitment to phagosomes. **A)** Representative Western blots of WT MEFs transfected with siCTR or siSec22b (50 nM) for 48 h incubated with anti-Sec22b (SYSY) and anti-tubulin as control (related to Figs. 5E and 6D). **B-C)** Representative Western blots of WT MEFs transfected with siCTR (100 nM) or siORP5/8 (50+50 nM) for 48 h incubated with anti-ORP8 **(B)** or and anti-ORP5 **(C)** and anti-tubulin as control (related to Fig. 5C). In (B) the two bands correspond roughly to the molecular weight of the canonical (100 kDa, closed arrow) and short (80 kDa) isoform (ORP8s, open arrow). In (C) the band corresponding to the typical running molecular weight of the canonical ORP5 isoform (110 kDa, closed arrow) responded to siRNA treatment while two other bands (presumed non-specific) did not. **D)** Quantification of Western blots sets represented in A-C shows efficient knockdown for Sec22b and canonical ORP isoforms (n=8;7;4;3 for siCTR;siSec22b;ORP8 in siORP5/8; ORP5 in siORP5/8). **E)** Confocal images of mCh-STIM1 and shR-EGFP-Sec22b-P33 in shSec22b MEFs exposed to IgG-beads for 30 min (related to Fig. 6F). Arrows: periphagosomal puncta reminiscent of MCS show co-localization of STIM1 and P33. White bar = 3 µm. Bar graphs are means + SEM. *p<0.05, **p<0.01.

**Figure S4.**
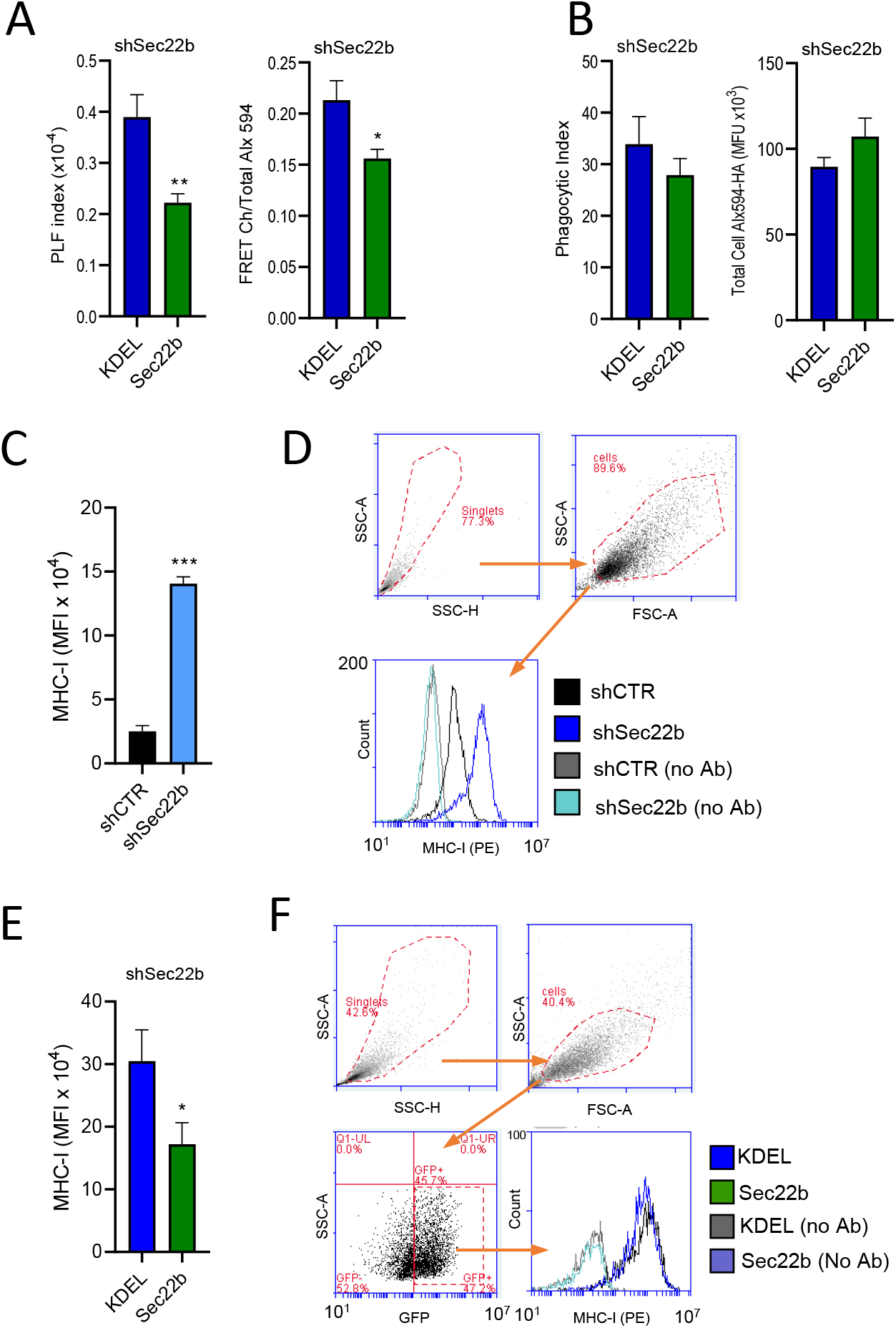
Related to Figure 7. Phago-lysosome fusion at high phagocytic loads and MHC-I cell surface expression. **A)** PLF index measured as in Fig. 6 in phagocytic shSec22b MEFs transfected with GFP-KDEL or shR-EGFP-Sec22b, except cells were exposed at 1:50 cells:AF488-IgG-bead ratio. Left panel: quantification of the PLF index as in Fig. 6. Right panel: Alternate calculation of PLF index (FRET channel normalized to total cell acceptor loading) that omits the GFP/AF488 donor channel normalization. In both cases PLF is decreased in cells expressing Sec22b as compared to KDEL controls. (n=6). **B)** Quantification of phagocytic index (left panel) and total cell loading of AF549-HA (right panel) of the cells analysed in (A). **C)** Mean fluorescence intensity (MFI) of cell surface staining of shCTR and shSec22b phagocytic MEFs using anti-H-2kb-PE, analysed by flow cytometry. Surface MHC-I was highly increased in shSec22b cells (n=4). **D)** Representative gating strategy of flow cytometry data in (C). **E)** Similar measurements as in (C) in shSec22b phagocytic MEFs expressing GFP-KDEL or shR-EGFP-Sec22b, gated on GFP+ cells. Surface MHC-I was reduced in cells transfected with Sec22b as compared to KDEL controls. (n=4). **F)** Representative gating strategy of flow cytometry data in (E).

